# Vimentin promotes actin assembly by stabilizing ATP-actin subunits at the barbed end

**DOI:** 10.1101/2025.11.17.688760

**Authors:** Lilian Paty, Lukas Kalvoda, Maritzaida Varela-Salgado, Quang D. Tran, Martin Lenz, Antoine Jégou, Guillaume Romet-Lemonne, Cécile Leduc

**Affiliations:** Université Paris Cité, CNRS, Institut Jacques Monod, Paris, France; Université Paris-Saclay, CNRS, Laboratoire de Physique Théorique et Modèles Statistiques, Orsay, France; Department of Physiology and Biomedical Engineering, Mayo Clinic, Rochester, MN 55905, USA; Physique et Mécanique des Milieux Hétérogènes (PMMH), CNRS, Ecole Supérieure de Physique et de Chimie lndustrielles de la Ville de Paris, Paris Science et Lettres Research University, Sorbonne Université, Université Paris Cité, Paris, France

**Keywords:** cytoskeleton, intermediate filaments, assembly dynamics, cytoskeletal crosstalk, in vitro reconstitution

## Abstract

Vimentin intermediate filaments play essential roles in maintaining cell integrity and regulating numerous cellular functions. In particular, vimentin cooperates with the actin cytoskeleton in key cellular processes that rely on actin dynamics, such as migration, division, and mechanosensing. While there is evidence that these two cytoskeletal components interact in cells, the underlying molecular mechanisms are only partially understood. Actin and vimentin can interact through biochemical signaling pathways or via cross-linkers, but whether they engage in a direct protein-protein interaction has remained controversial, in part because such interactions are difficult to isolate and characterize in cells. Using in vitro reconstitution coupled to theoretical modeling, and total internal reflection fluorescence microscopy to monitor the elongation of single actin filaments, we show that vimentin promotes actin elongation by stabilizing actin subunits at the barbed end in a dose-dependent manner. Strikingly, this effect depends on the nucleotide state of actin, as the acceleration is only observed for the elongation from ATP-actin, and not ADP-actin monomers. We further establish that neither the vimentin tail nor head domains are required for this effect, and both filamentous and non-filamentous vimentin enhance actin elongation. Finally, we find that vimentin promotes the nucleation of actin filaments. Consistently, magnetic pull-down assays demonstrate a direct interaction between vimentin and ATP-actin monomers. Altogether, these findings identify vimentin as an unexpected new actor in the regulation of actin dynamics at the barbed end and bring new insights into the functional role of vimentin through cytoskeletal crosstalk.

## Introduction

The cytoskeleton is a dynamic network of three interconnected filament systems: actin filaments, microtubules, and intermediate filaments, that allow cells to shape and organize themselves in space and time while mechanically engaging with their environment. Intermediate filaments are composed of a large family of proteins whose expression levels are cell- and tissue-type specific (1–4). Among them, vimentin is expressed in mesenchymal cells and widely used as a marker of the epithelial-to-mesenchymal transition, during which epithelial cells acquire migratory and invasive properties (5, 6). Unlike actin and microtubules, which rapidly remodel through active turnover, vimentin self-assembles into long-lived apolar filaments (3, 7–9), yet is involved in highly dynamic processes such as migration, division, and mechanosensing (4, 10–13). Growing evidence suggests that vimentin contributes to these functions through its crosstalk with the other cytoskeletal filaments (2, 14–16), including actin (17). Early studies showed that vimentin-deficient cells exhibit reduced acto-myosin contractility (12). More recent work revealed that vimentin coordinates actin stress fiber assembly (18–20), orients traction forces that drive migration (21), and tunes cell stress by assisting in actomyosin-based force transmission (10). There is also a mechanical synergy between the two types of filaments (22), which is crucial for maintaining cell integrity (23). The actin-vimentin interplay likely relies on reciprocal regulation of filament organization and dynamics, with vimentin modulating actin in both non-dividing (18, 24–26) and dividing cells (11, 13) and actin in turn influencing vimentin (18, 27–29). This interplay is mediated through indirect biochemical signaling, direct interactions, or interactions via crosslinkers (2, 14, 18, 30-33), but the precise molecular mechanisms remain to be fully elucidated.

In vitro reconstitution with purified proteins provides a powerful approach to dissect how actin and vimentin coordinate their activities, as it can disentangle their interactions from the complexity of the cellular environment. Bulk rheology studies of actin-vimentin networks have suggested that the observed viscoelastic properties of the composite networks cannot be explained without direct actin-vimentin interactions (34, 35), and that vimentin’s tail domain is implicated in the interaction (34). However, another study has argued that the mechanics of composite networks could be explained solely by the additive properties of their individual components, without requiring direct filament-filament binding (36). More recently, single-filament imaging found no evidence for actin-vimentin filament co-alignment, suggesting that if there is an interaction between the filaments, it must be transient and weak (37). Thus, whether actin and vimentin interact directly, and how such interactions might influence filament dynamics, remains unresolved. Understanding the effect of vimentin on actin filaments alone in simplified in vitro systems is necessary, before adding more components, such as crosslinkers and regulatory proteins, in more complex bottom-up approaches that would come closer to the cellular context.

Here, using an actin single-filament assay coupled to theoretical modeling, we found that vimentin directly promotes actin filament nucleation and elongation. We show that the molecular mechanism responsible for the enhanced elongation involves vimentin stabilizing actin subunits at the barbed end in a nucleotide state-dependent manner. These findings reveal a previously unrecognized role for vimentin as a regulator of actin filament growth, highlighting a direct interaction through which vimentin intermediate filaments can influence actin dynamics.

## Results

### Vimentin promotes actin filament elongation at the barbed end

In order to assess the impact of vimentin on actin assembly dynamics, we used an assay that allows direct visualization of actin elongation at the single filament level using time-lapse Total Internal Reflection Fluorescence (TIRF) microscopy, hereinafter referred to as “actin polymerization assay” (Fig. 1A). Small pre-polymerized biotinylated fragments of actin filaments (F-actin) (Fig. 1A, magenta) were first anchored to a passivated glass surface coated with neutravidin. Upon addition of 0.5 µM of actin monomers in the ATP-bound state (ATP-G-actin, 10% fluorescently labeled), with a low fraction of biotinylation (∼0.15%), filaments elongated at their fast-growing barbed-end with sparse anchoring points along their length (∼0.5 anchors per µm) (Fig. 1A-B, yellow). At this actin concentration, the elongation of the pointed end is negligible over the time scale of the experiments. This low biotinylation level minimized potential effects on elongation while maintaining sufficient proximity to the surface for automated filament tracking. Actin filament elongation was monitored in either assembly buffer alone (control) or buffer containing pre-assembled vimentin filaments at 1 mg/ml (18 µM vimentin monomers). Automated filament tracking and systematic quantification of the elongation rate (Fig. 1C) demonstrated that actin filaments elongated faster in the presence of vimentin filaments than in the control condition (Fig. 1D). This effect was consistently reproducible across repeated experiments (N = 11 independent replicates), and robust with salinity (Fig. S1). Control experiments verified that the actin elongation rates with low biotinylation were comparable to those without anchoring points, using methylcellulose to confine the filaments (Fig. S2A). Moreover, we verified that vimentin’s effect on actin elongation was similar in both anchoring and methylcellulose conditions (Fig. S2B). However, because methylcellulose induces actin-vimentin co-alignment (potentially bundling them through depletion effects), as observed when a small fraction of vimentin was fluorescently labeled within the network (Fig. S2C), we rather used sparse anchoring to confine the filaments on the surface. Because, in cells, actin monomers are bound to profilin and F-actin elongate from profilin-G-actin complexes, we verified that vimentin also promoted elongation from profilin-G-actin (Fig. S3). These results demonstrate that the sole presence of vimentin filaments is sufficient to accelerate actin elongation in the absence of any actin-vimentin crosslinkers.

**Figure 1:**
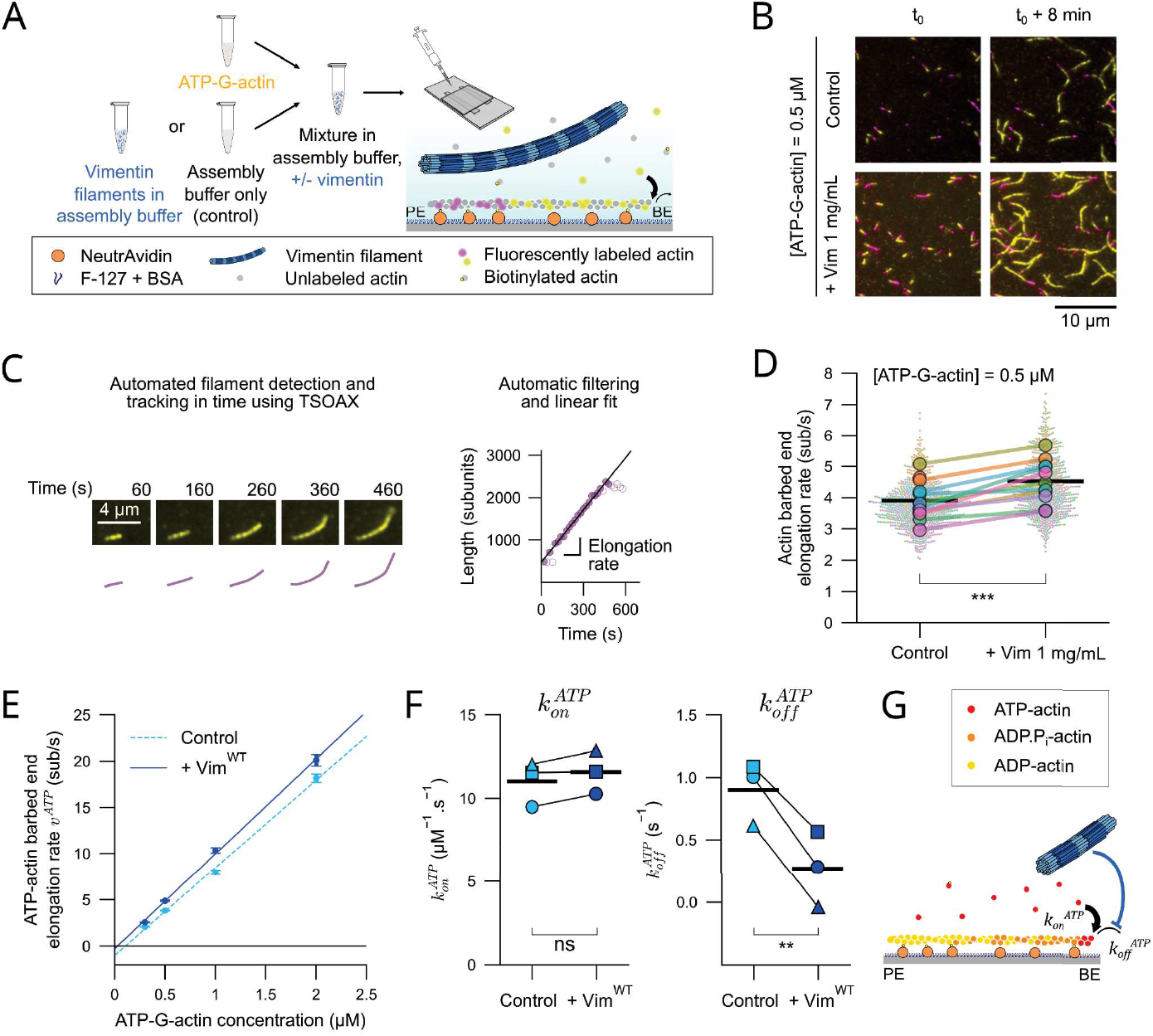
Vimentin promotes actin filament elongation at the barbed end by stabilizing ATP-actin subunits. A) Actin polymerization assay. Fragments of actin filaments (10% Atto-643, 10% biotin labeled, magenta) were pre-polymerized for >1h at 10 pM by adding 100 mM KCI to the ATP-G-actin solution in G-buffer, then diluted and fragmented by pipetting and introduced in a flow chamber that was functionalized with neutrAvidin and passivated by F127 and BSA. ATP-G-actin (10% AF-488 or Atto-488, 0.15% biotin labeled, yellow) was mixed at a concentration of 0.5 pM in assembly buffer (10 mM HEPES pH 7.0, 1 mM sodium phosphate, 1 mM MgCI2, 0.2 mM EGTA, 100 mM KCI) only (Control) or in buffer with 1 mg/mL of pre-assembled unlabeled wild-type vimentin filaments. This solution was then introduced in the flow chamber, and actin monomers assembled from the barbed end of the anchored F-actin fragments in the presence or absence of vimentin (blue). Actin filament elongation was monitored by TIRF microscopy. PE: pointed end, BE: barbed end. B) Representative TIRF snapshots of actin filaments polymerizing without (Control) or with (+ Vim 1 mg/mL) vimentin filaments C) Method used to quantify the actin barbed end elongation rate. Left: example of TIRF time-lapse sequence of an elongating filament, along with the corresponding (68)entation by the software TSOAX (66). Right: length in subunits of this actin filament as a function of time. The elongation rate was quantified by automatic filtering of detection errors (open circles) and linear fitting. Detection errors are due to the sparse anchoring of the filament: between two anchoring points, the filament can sometimes grow out-of-plane, leading to a low TIRF signal and an underestimation of its length. D) Actin barbed end elongation rate (subunits per second) in the absence (Control) or in the presence (+ Vim 1 mg/ml) of vimentin filaments. Colors depict experiments performed on the same day. Small dots: individual actin filament elongation rates *(n* ≥ 20 rates per experiment), large dots: average elongation rates of the replicates *(N* = 11 independent replicates), black bars: mean elongation rate computed from the replicate averages. Statistical significance of the difference was evaluated by running a two-tailed paired t-test over the replicate averages. ***, p<0.001. E) Actin barbed end elongation rate as a function of ATP-G-actin concentration in the absence (Control, light blue) or in the presence of vimentin filaments at 1 mg/ml(+ Vimwr, dark blue). Data are shown as the mean ± standard error of the mean for *n ≥*20 rates per condition.Dashed and solid lines represent linear fits of the means to the equation 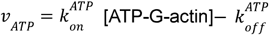, to extract the on- and off-rate constants 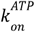and 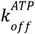.Experiments were conducted in the standard assembly buffer at 50 mM KCI. F) Quantification of the ATP-actin on- and off-rate constants at the barbed end 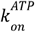and 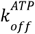 extracted from *N* = 3 independent replicates (filled symbols) and mean rates (black bars), in the absence (Control, light blue) or in the presence of vimentin filaments (+Vim^WT^, dark blue) at 1 mg/ml. Statistical significance of the difference between conditions was evaluated by running two-tailed paired t-tests over the replicate averages. **, p<0.01, ns: not significant (p>0.05). G) Schematics of the nucleotide states of actin in the standard actin polymerization assay. The barbed end (BE) is occupied by ATP-actin subunits (red), and free ATP-actin monomers in solution bind and unbind from the barbed end with rate constants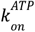 and 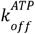,respectively. Deeper within the filament, actin subunits exist as ADP.P;-actin (orange) and ADP-actin (yellow). Vimentin (blue) increases the net ATP-actin elongation rate at the barbed end by decreasing the ATP-actin off-rate constant 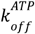.

### Vimentin promotes actin elongation by preventing the departure of ATP-actin subunits

The net elongation rate of actin filaments at the barbed end reflects the result of actin monomer association and dissociation events. To gain insights into the mechanism by which vimentin accelerates actin elongation, we sought to determine how the arrival and the departure of monomers at the barbed end were respectively affected by vimentin. To do so, we repeated the actin polymerization assay at different ATP-G-actin concentrations (Fig. 1E). The elongation rates displayed an affine relationship with the ATP-G-actin concentration, allowing the extraction of the on-rate (slope) and the off-rate (intercept) constants, called 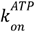and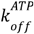 respectively. Strikingly, we observed that vimentin reduced the actin off-rate constant close to zero, while having no significant effect on the on-rate constant (Fig. 1F). There may be a minor effect on the on-rate, but the dominant effect is on the off-rate. Thus, vimentin accelerates actin elongation by stabilizing monomers at the growing barbed end rather than increasing the monomer binding rate (Fig. 1G).

### Vimentin does not stabilize actin filaments against depolymerization

The stabilization of ATP-actin subunits at the barbed end induced by vimentin during polymerization led us to test whether vimentin could also protect actin filaments from depolymerization. We performed an actin depolymerization assay where non-biotinylated actin filaments were first polymerized from surface-anchored spectrin-actin seeds. Then, unpolymerized actin monomers were washed out, and filament depolymerization was monitored using methylcellulose to keep the filaments close to the surface (Fig. 2A). The presence of vimentin did not decrease the actin depolymerization rate, indicating that vimentin stabilizes the barbed end only when the actin filament is growing (Fig. 2A). On the contrary, we observed a mild increase in the depolymerization rate. This surprising result might be explained by the difference in state of the barbed end under polymerizing and depolymerizing conditions. Upon incorporation into a filament, the conformational change of the subunits triggers the ATP hydrolysis to ADP.Pi within seconds, and the inorganic phosphate is then released with a half-time of a few minutes (38–40). Actin subunits can thus exist under three different nucleotide states (ATP-, ADP.Pi- and ADP-actin). In polymerizing conditions, when the polymerization rate is faster than the ATP hydrolysis rate (0.3 s-^1^ (41)), the barbed end is mostly occupied by ATP-actin subunits, and the off-rate measured indirectly corresponds to the dissociation of ATP-actin (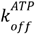 subunits Fig. 1G). In contrast, in the depolymerization assay, we monitor the disassembly of ADP/ADP.Pi-actin subunits(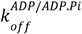 Fig. 2A). Note that we do not monitor the depolymerization of fully ADP-actin filaments, as the buffer contains 1 mM sodium phosphate, which comes with vimentin, preventing the full conversion to ADP-actin filaments by maintaining an equilibrium between filament-bound and soluble phosphate (see Methods). The fact that vimentin has no impact on ADP/ADP.Pi depolymerization suggests that its stabilizing action at the barbed end is specific to ATP-actin subunits.

**Figure 2:**
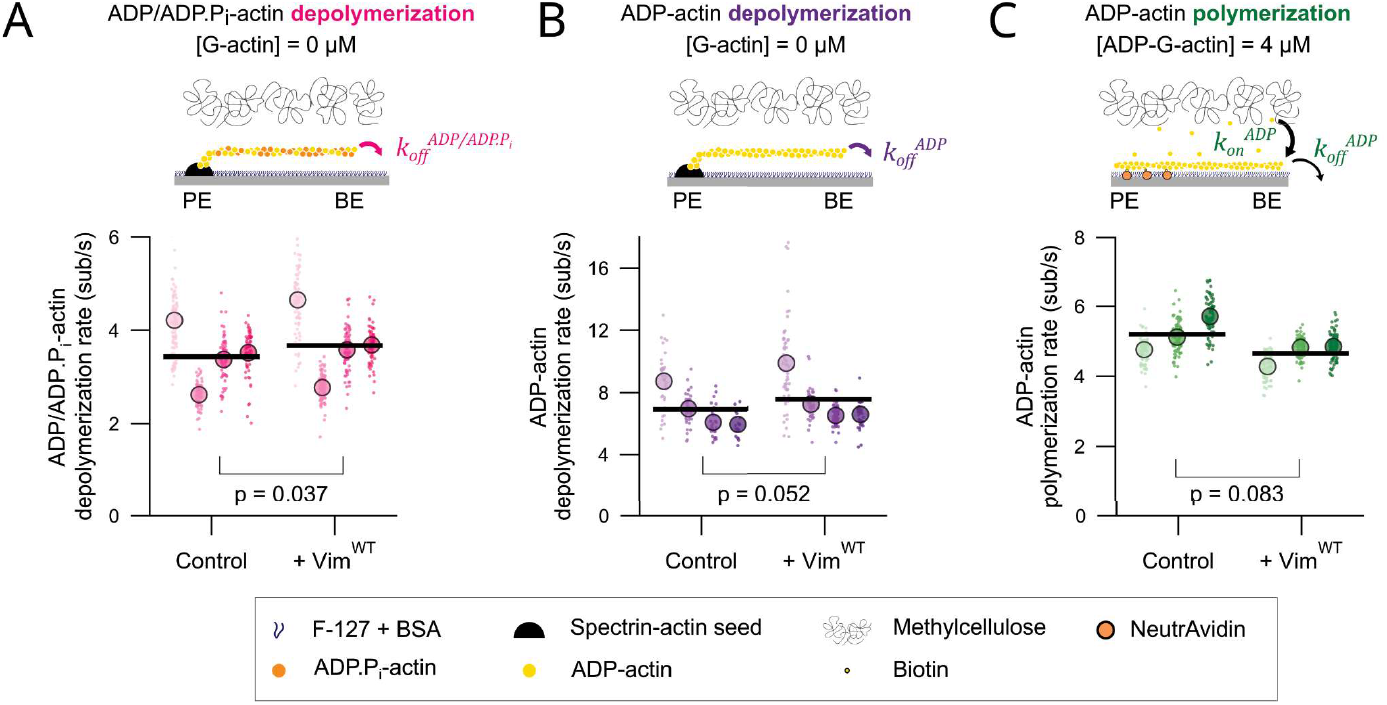
Vimentin’s stabilizing effect at the barbed end is nucleotide-dependent. A) Top: Actin depolymerization assay in Tris assembly buffer (10 mM Tris-HCI, pH 7.0, 1 mM MgCl_2_, 0.2 mM EGTA, 50 mM KCI). Actin filaments were polymerized from spectrin-actin seeds by flowing ATP-actin monomers in the flow chamber. Non-polymerized actin monomers were then rinsed. After waiting for five minutes for ATP to get hydrolysed and inorganic phosphate to be released, the depolymerization of actin filaments was monitored by TIRF microscopy, keeping the filaments close to the surface using methylcellulose. Bottom: ADP-actin depolymerization rate in the absence (Control) or in the presence (+ Vim^WT^) of vimentin filaments at 1 mg/ml. Small dots: individual depolymerization rates (n 2≥ 20 rates per replicate), large dots: average depolymerization rates of the replicates *(N* = 4 independent replicates), black bars: mean depolymerization rate computed from the replicate averages. B) Top: ADP-actin polymerization assay in Tris assembly buffer. ADP-G-actin was obtained by incubating ATP-G-actin at 10 μM in ATP-free G-buffer supplemented with 2.5 mM glucose, 0.2 mM EGTA, 20 μM MgCl_2_ and 15 U/ml hexokinase for 18 h on ice. Unlabeled ADP-G-actin was mixed with 50 nM Alexa488-labeled actin-binding domain of utrophin in Tris assembly buffer and introduced in a flow chamber to polymerize from anchored F-actin fragments. The ADP-G-actin concentration was 4 μM. The filaments were kept close to the surface by methylcellulose and their elongation was followed by TIRF microscopy. Bottom: ADP-actin polymerization rate in the absence (Control) or in the presence (+ Vim^WT^) of vimentin filaments at 1 mg/ml. Small dots: individual polymerization rates (n 2≥20 rates per replicate), large dots: average polymerization rates of the replicates *(N* = 3 independent replicates), black bars: mean polymerization rate computed from the replicate averages. Statistical significance of the differences between conditions in A, B and C was evaluated by running two-tailed paired t-tests over the replicate averages. In A, Band C, color shades denote paired and comparable experiments. C) Top: ADP-actin polymerization assay in Tris assembly buffer. ADP-G-actin was obtained by incubating ATP-G-actin at 10 μM in ATP-free G-buffer supplemented with 2.5 mM glucose, 0.2 mM EGTA, 20 μM MgCl_2_ and 15 U/ml hexokinase for 18 h on ice. Unlabeled ADP-G-actin was mixed with 50 nM Alexa488-labeled actin-binding domain of utrophin in Tris assembly buffer and introduced in a flow chamber to polymerize from anchored F-actin fragments. The ADP-G-actin concentration was 4 μM. The filaments were kept close to the surface by methylcellulose and their elongation was followed by TIRF microscopy. Bottom: ADP-actin polymerization rate in the absence (Control) or in the presence (+ Vim^WT^) of vimentin filaments at 1 mg/ml. Small dots: individual polymerization rates (n 2≥20 rates per replicate), large dots: average polymerization rates of the replicates *(N* = 3 independent replicates), black bars: mean polymerization rate computed from the replicate averages. Statistical significance of the differences between conditions in A, B and C was evaluated by running two-tailed paired t-tests over the replicate averages. In A, Band C, color shades denote paired and comparable experiments.

### Vimentin stabilizes ATP-actin, but not ADP-actin monomers at the barbed end

To assess the influence of the actin nucleotide state on the stabilizing effect of vimentin, we first repeated the depolymerization assay in a Tris buffer without phosphate (Fig. 2B). This allowed us to probe the impact of vimentin on the ADP-actin off-rate constant 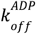. As observed in the phosphate-containing buffer, vimentin did not have a stabilizing effect on the barbed end, suggesting that vimentin should not be able to accelerate actin filament elongation from ADP-G-actin. To verify this, we monitored actin polymerization from ADP-actin monomers (ADP-G-actin) by converting ATP-G-actin into ADP-G-actin prior to the assay (Fig. 2C). Because ADP-actin has a higher critical concentration than ATP-actin (38, 42, 43), we performed the actin polymerization assay at 4 µM ADP-G-actin instead of 0.5 µM, and in a Tris buffer without phosphate to avoid any conversion to ADP.Pi. However, this higher actin concentration increased the background signal when a small fraction of actin was fluorescently labeled, preventing accurate tracking of filament elongation. To overcome this constraint, we used a low concentration (50 nM) of a fluorescently labeled actin filament–binding domain from utrophin, which reliably reports filament length without interfering with assembly (44, 45) or with the effect of vimentin on actin elongation (Fig. S4). Unlike with ATP-actin, we did not observe any acceleration of ADP-actin polymerization by vimentin, confirming that the stabilizing effect of vimentin depends on the nucleotide state of the actin subunits and occurs only for ATP-actin. Taken together, these results demonstrate that vimentin can stabilize the barbed end of actin filaments when occupied by ATP-actin subunits, thereby enhancing polymerization. In contrast, when the barbed end is composed of ADP-actin subunits, vimentin has no stabilization effect, which explains the absence of an effect on actin depolymerization.

### Non-filamentous vimentin also promotes actin elongation

We next asked whether vimentin’s effect on actin elongation depends on vimentin’s filament-forming capacity. In vimentin-expressing cells, vimentin is found as elongated filaments, short filaments also called squiggles, and tetramers (4, 46, 47). At low ionic strength, purified vimentin spontaneously self-assembles into tetramers (3, 48). By raising the ionic strength, tetramers associate laterally into short filaments known as unit-length filaments (ULFs), which subsequently anneal end-to-end to form longer filaments (3, 48). To arrest assembly at different stages, we produced different mutants of vimentin (Fig. 3A). The headless mutant (Vim^Δhead^), which lacks the N-terminal disordered head domain, forms tetramers but fails to laterally assemble into ULFs (48). The point-mutated vimentin Vim^Y117L^ forms ULFs that cannot anneal end-to-end to form longer filaments (49). Performing the actin polymerization assay (Fig. 1A) with these vimentin mutants revealed the same increase in actin elongation rate as observed with wild-type vimentin (Vim^wt^), compared with the vimentin-free control condition (Fig. 3B). We conclude that the effect of vimentin on actin elongation can occur even with vimentin tetramers, and does not require vimentin to be assembled into long filaments.

**Figure 3:**
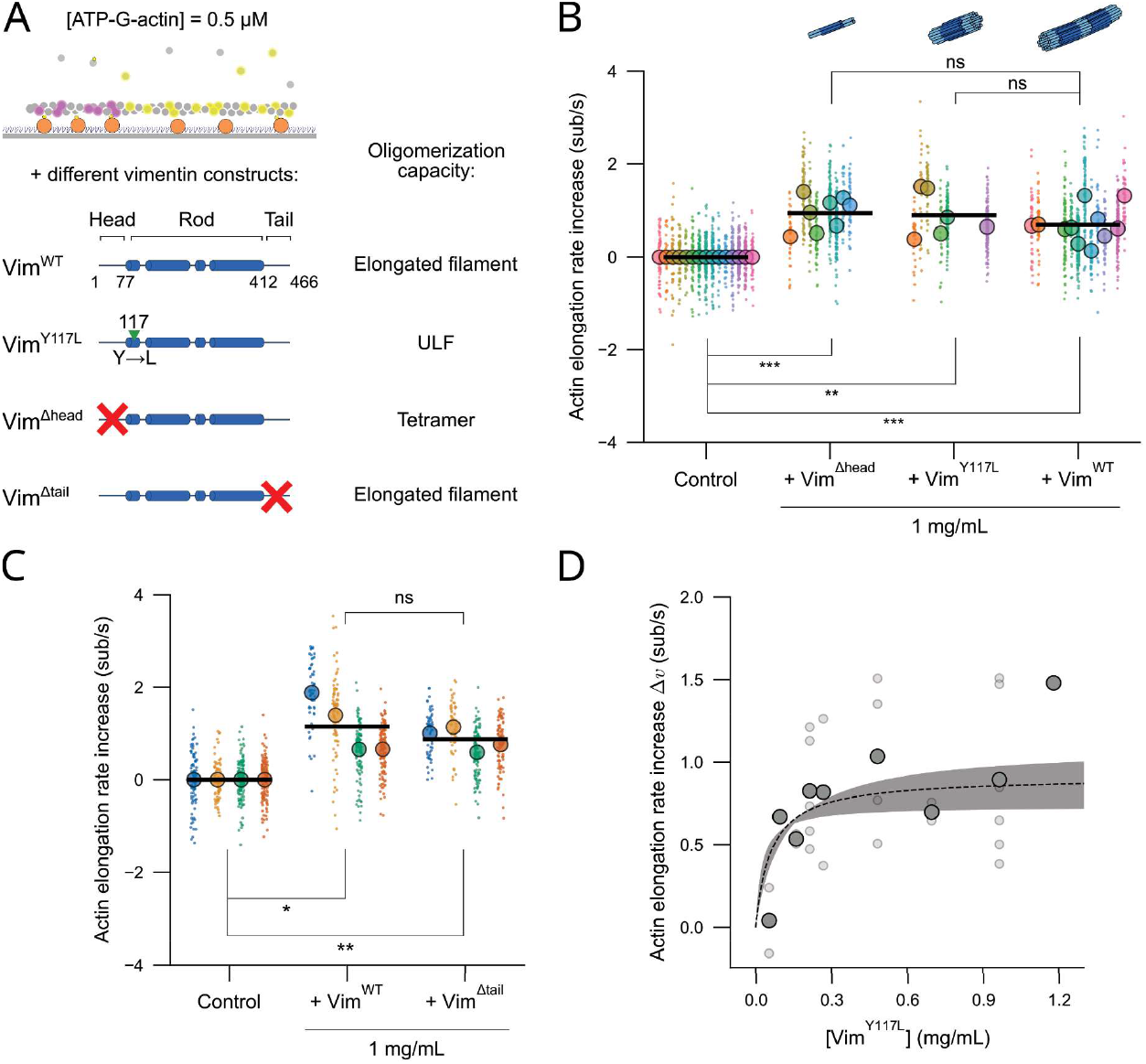
Vimentin acceleration of actin elongation is independent of its oligomerization state and tail domain, and is dose-dependent. A) The actin polymerization assay (at 0.5 μM ATP-G-actin, 10% labeled) was performed in the presence of different vimentin constructs. Wild-type vimentin (Vim^WT^) and tailless vimentin (Vim^Δtail^) assemble into filaments. Vim^Y117^Lassembles into unit-length-filaments (UlFs) that do not anneal end-to-end. Headless-vimentin (Vim^Δhead^)forms tetramers that do not assemble into filaments. The numbers represent the positions of the residues that delimit the head and tail domains. B) Actin elongation rate increase in the presence of 1 mg/ml vimentin (18 μM vimentin monomers) under different oligomerization states (+ Vim^Δhead^, VimY^Y117L^ or Vim^WT^), compared to the average rate of the control condition for each replicate. Colors depict experiments performed on the same day. Small dots: individual elongation rate increases *(n* ≥ 20 rates per replicate), large dots: average elongation rate increases of the replicates (Vim^Δhead^ : *N* = 8, Vim^Y117L^ : *N* = 6, Vim^WT^: *N* = 11 independent replicates), black bars: mean elongation rate increase computed from the replicate averages. The data shown for wild-type vimentin are the same as shown in Fig. 1D. C) Actin elongation rate increase in the presence of 1 mg/ml of Vim^WT^ or Vim^Δtail^ filaments. Colors depict experiments performed on the same day. Small dots: individual elongation rate increases *(n* ≥ 20 rates per replicate), large dots: average normalized elongation rates of the replicates *(N* = 4 independent replicates), black bars: mean elongation rate increase computed from the replicate averages. Experiments in C were conducted in the standard assembly buffer at 50 mM KCI. Statistical significance of the differences between conditions in B and C was evaluated by running two-tailed paired t-tests over the replicate averages. * p<0.05, ** p<0.01, *** p<0.001, ns: not significant (p>0.05). D) Actin elongation rate increase *(Δ v)*, compared to the average rate of the control condition (without Vim^Y117L^) measured on the same day, as a function of Vimv^Y117L^ concentration (*[Vim*^Y117L^]). 1 ≤*N≤* 8. Small dots: average rate increase of individual replicates *(n*≥ 20 rates per replicate and condition), large dots: average of the replicates for each concentration (1≤*N*≤8 replicates per concentration). Dashed line: fit of the unbinned data with the binding model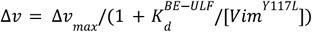, where the only unknown parameter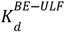 characterizes the putative interaction between the actin barbed end and Vim^Y117L^ ULFs (see Theoretical modeling in SI). The plateau value *dv* is fixed to its mean value Δ*v*_*max*_ obtained from the ATP-G-actin on- and off-rate constants of the *N* = 3 replicates in Fig. 1F. The fit yields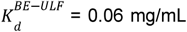. Shaded area: error in the fitted elongation rate increaseΔ*v* due to uncertainties in its plateau Δ*v*_*max*_. We vary the plateau value Δ*v*_*max*_ within the range given by its standard error (SEM). This leads to a continuum of fitted curves, with values of 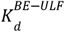 in the range (0.02 - 0.11) mg/ml.

### Vimentin’s disordered tail domain is not required to promote actin elongation

Because vimentin’s disordered tail domain has been proposed to mediate a direct actin-vimentin interaction (34), we asked whether this domain is required for the acceleration of actin assembly. To address this question, we produced a truncated version of vimentin lacking the last 55 amino acids (Vim^Δtail^) and performed the actin polymerization assay (Fig. 1A) in the presence of this mutant. This tailless vimentin mutant assembles into filaments (3, 48). Vim^Δtail^ increased the actin elongation rate to the same extent as wild-type vimentin, with no significant difference between the two conditions (Fig. 3C), indicating that the tail domain is not required to promote actin elongation at the barbed end.

### Vimentin promotes actin filament elongation in a dose-dependent manner

We next investigated whether vimentin’s effect depends on the vimentin concentration. To this end, we took advantage of the vimentin Y117L mutant, which allows for the precise control of both the number and oligomerization state of vimentin (ULFs). Actin polymerization assays were performed with 0.5 µM ATP-G-actin in the presence of Vimv^117L^ ULFs at concentrations ranging from 0 to 1.18 mg/ml. The actin elongation rate difference relative to the control (Δ*v*) increased with the vimentin Y117L concentration ([Vim ^*Y117L*^]) (Fig. 3D). As a first simple model, we propose that the saturation stems from a putative interaction between vimentin Y117L ULFs and the actin barbed end. In this framework, the barbed end of a growing actin filament is in a dynamic equilibrium between two states: free or occupied by a vimentin ULF (see Theoretical modeling in SI). Accordingly, our polymerization assay probes the ATP-G-actin on- and off-rate constants in the two extreme cases of this model where the barbed end is either never (Control) or always occupied (+Vim), implying that the effect saturates at a corresponding maximum value Δ*v*_*max*_. Thus, we model the increase in actin elongation rate by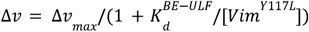, where the only unknown parameter 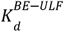 characterizes the global interaction between actin barbed ends and vimentin ULFs, and encompasses unresolved molecular details. Fitting this model to the dose-dependent acceleration of actin elongation by vimentin Y117L yields a dissociation constant of about 0.06 mg/ml. Hence, the effect of vimentin on actin elongation reaches saturation with only a limited amount of vimentin, as compared to the physiological range (estimated to be > 1 mg/ml in Hela cells (50)).

### Vimentin promotes actin nucleation

Vimentin’s effect on ATP-actin subunits at the barbed end significantly lowers the critical concentration for net ATP-actin elongation, defined as the ratio between the actin off-rate 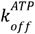 and the on-rate 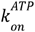 We hypothesized that vimentin could also enhance the nucleation of actin filaments, which requires the formation of a critical nucleus, most commonly described as a tetramer, although some models suggest a trimer (51–53). Because small oligomers are unstable, their formation represents the principal barrier to nucleation. By stabilizing the interaction between ATP-actin subunits, vimentin could reduce this barrier and promote actin oligomerization to form a critical nucleus. To test this hypothesis, we first analyzed the ability of vimentin to nucleate actin filaments in the bulk using a pyrene fluorescence assay (Fig. 4A). The presence of vimentin decreased the initial lag time associated with actin filament nucleation, and fitting the polymerization curves at early times (see Materials and methods in SI) using the off-rate 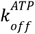 and on-rate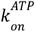 constants extracted from previous experiments (Fig. 1F) revealed a ∼35% increase in the initial nucleation rate (Fig. 4B, left). The differences in plateau intensity were also qualitatively consistent with the difference in critical elongation concentration we characterized. Fitting the slope of the polymerization curve at each time point further indicated that the actin barbed end concentration remains enhanced in the plateau regime (Fig. 4C). To support our empirical analysis, we propose a minimal theoretical model of F-actin nucleation and elongation subject to a limited monomer pool c_0_ = [*ATP* - *G* - *actin*](*t* = 0) (see Theoretical modeling in SI). We consider the nucleation of critical tetrameric F-actin nuclei with a rate constant *k*_*nucl*_, and ATP-G-actin depletion due to polymerization with a rate constant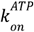 We reason that short filaments tend to disassemble when the free ATP-G-actin concentration c is below the critical concentration for elongation 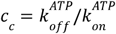As a result, nucleation events in this regime are typically unproductive and need not be considered in our formalism. We thus adopt an expression for the net nucleation rate *r*_*nucl*_(*c*) that vanishes at *c = c_c_*, and satisfies the *c*^4^ dependence at large actin concentrations consistent with tetrameric F-actin nuclei, 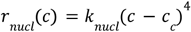 This allows us to predict the number of actin barbed ends per unit volume as a function of time

**Figure 4:**
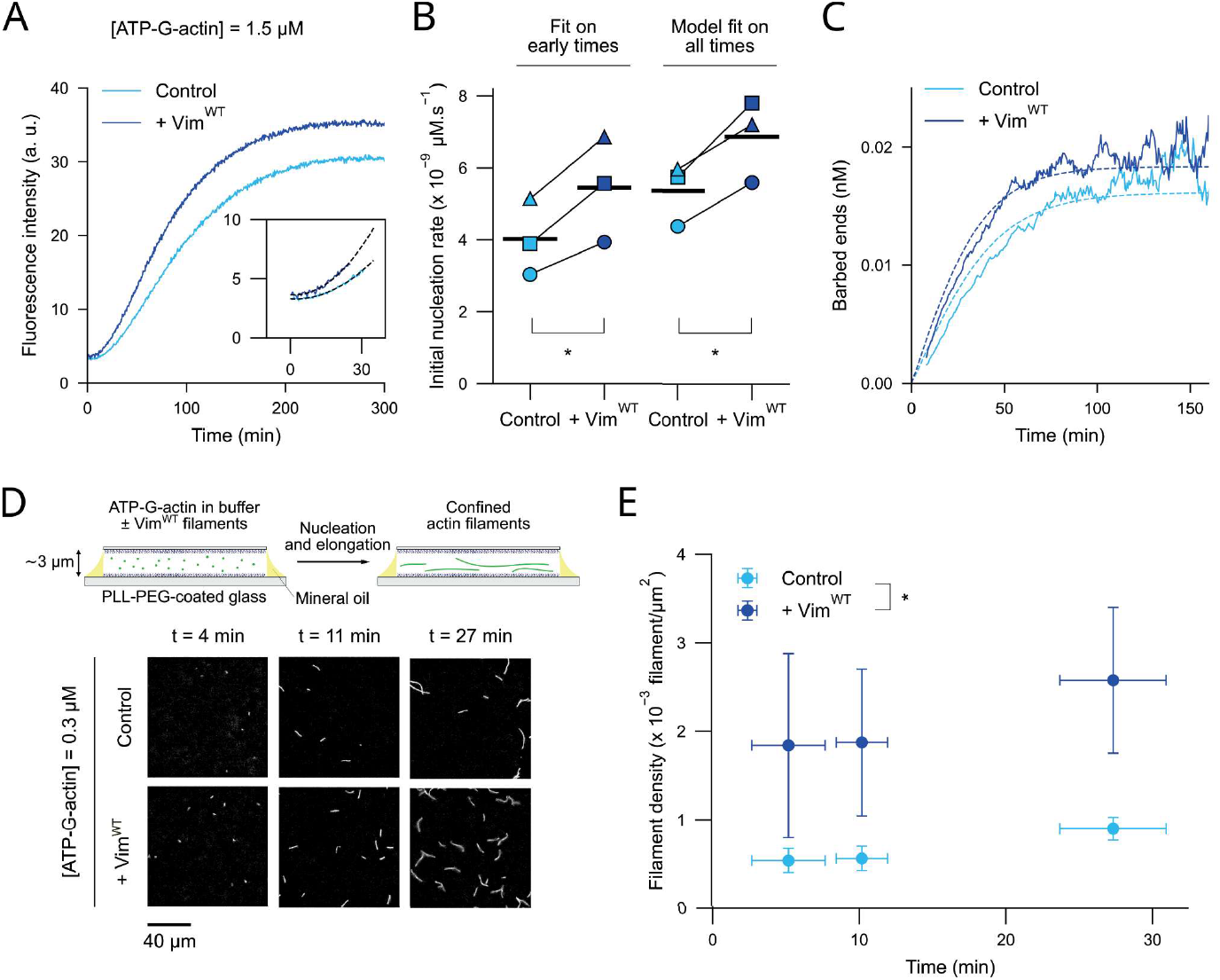
Vimentin promotes actin nucleation. A) Representative pyrene-actin fluorescence assay of actin assembly. Actin (1.5 μM, 8% pyrene-labeled) was assembled in the absence (Control, light blue) and in the presence of 1 mg/ml of pre-assembled wild-type vimentin filaments (+ Vim^WT^, dark blue). Raw fluorescence signals were corrected for the vimentin signal (see Materials and methods in SI). The inset shows an enlarged plot of the early time points, which were first corrected for photobleaching (Fig. S5A) and then fitted based on previously described method (69) to extract the initial nucleation rate (black dashed lines, see Materials and methods in SI). This representative replicate corresponds to the one represented with squares in the quantification in B. B) Quantification of the initial nucleation rate for N = 3 independent replicates in control (light blue) and vimentin (+ Vim^WT^, dark blue) conditions, using two different methods: empirical one-parameter fit on early time points as previously described (69)(Ieft), or one-parameter fit of the whole polymerization curve using our theoretical model (right, Fig. S5B). Symbols: initial nucleation rates measured for each replicate with both methods, black bars: mean initial nucleation rates over the 3 replicates. Statistical significance of the difference between conditions was evaluated by running two-tailed paired t-tests. * p<0.05. C) Concentration of actin barbed ends over time in the absence (Control, light blue) and in the presence of 1 mg/ml of pre-assembled wild-type vimentin filaments (+ Vim^WT^, dark blue), for the replicate shown in A. Plain curves: barbed end concentration computed from the rate of actin polymerization over time, given by the slope at each time point along the fluorescence curve ((53), see Materials and methods in SI). Dashed lines: barbed end concentration predicted by our theoretical model described in the main text, following a one-parameter fit of the polymerized actin concentration over time (Fig. S5B and Theoretical modeling in SI). D) Direct visualization of actin nucleation by epifluorescence microscopy. Top: schematic representation of the experiment. ATP-G-actin (0.3 μM, 10% labeled) was mixed in assembly buffer with or without pre-assembled wild-type vimentin filaments, and a small volume of the sample was confined between two passivated glass coverslips, resulting in a ∼3-μm-high chamber. The chamber was sealed using mineral oil. Bottom: representative snapshots of nucleated actin filaments in the absence (Control) or in the presence of 1 mg/ml of pre-assembled wild-type vimentin filaments(+ Vim^WT^). E) Quantification of the mean filament density in C) as a function of time in control (light blue) and vimentin (+ Vim^WT^, dark blue) conditions. Each dot represents the mean+/- SEM of N = 4 replicates for a time interval depicted by the horizontal error bar. For each replicate, condition, and time point, the mean number of filaments from 24≤n≤94 fields of view was computed. Statistical significance of the difference between conditions was evaluated by running a two-way ANOVA. * p<0.05.

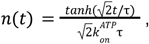

where we have defined the time scale 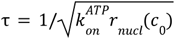 over which ATP-G-actin is depleted. The only unknown parameter, namely the nucleation rate constant *k*_*nucl*_, is fitted from the actin polymerization curve (Fig. S5B), which allows us to compute the initial nucleation rate *r*_*nucl*_ (*c*_0_), confirming that this rate is increased by vimentin (Fig. 4B, right). The quantitative mismatch with the empirical analysis may result from assuming the same critical concentration for nucleation as for elongation. Also, the model does not take into account experimental variability in the actin on- and off-rates and how vimentin affects them. Nevertheless, the good agreement between the model and the empirical analysis in the number of barbed ends over time (Fig. 4C) corroborates that vimentin enhances both actin nucleation and elongation. We then designed a fluorescence microscopy assay to directly visualize the nucleation of actin filaments. We mixed ATP-G-actin at low concentration (300 nM, 10% fluorescently labeled) in assembly buffer without or with pre-formed vimentin filaments, and confined the sample between two passivated glass coverslips, allowing us to detect and count filaments over time (Fig. 4D). We observed that the addition of vimentin filaments increased the number of actin filaments created by unit of time (Fig. 4E). Taken together, these results show that vimentin, beyond lowering the actin critical concentration for elongation, also accelerates filament formation, providing a direct mechanism by which vimentin intermediate filaments can modulate actin network assembly.

### Vimentin interacts directly with actin monomers

The effect of vimentin on actin nucleation suggests the existence of a direct interaction between vimentin and actin monomers. To test this, we designed a pull-down assay in which varying amounts of vimentin wild-type filaments were immobilized on magnetic beads and used to capture ATP-G-actin (Fig. 5A). Pellets and supernatants were analyzed by SDS-PAGE followed by gel staining, with known concentrations of actin and vimentin loaded on the same gel to generate calibration curves (Fig. 5B). We observed that the fraction of actin recovered in the pellet increased with the amount of vimentin on the beads (Fig. 5C). We fitted the fraction of actin specifically bound to the vimentin-coated beads *(p)* as a function of the free vimentin concentration in the pellet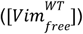 with the equation: 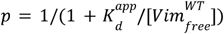, where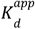 is the apparent dissociation constant between filamentous vimentin and ATP-G-actin, when assuming a 1:1 binding stoichiometry between ATP-G-actin and vimentin monomers (Fig. 5D, Theoretical modeling in SI). Pooling the data from 5 independent experiments allowed us to estimate 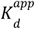to be of the order of 0.3 mg/ml, corresponding to 6 µM vimentin monomers. This value should be considered approximate. First, actin in solution may not be strictly monomeric due to spontaneous nucleation, potentially leading to an underestimation of the apparent dissociation constant. Second, although our quantification method accounts for the non-specific binding of actin to the beads, such binding may be reduced when beads are coated with vimentin, which would instead lead to an overestimation of the apparent dissociation constant. As we do not resolve the molecular details of the interaction, we analyzed the effect of different stoichiometries (Fig. S6 and Theoretical modeling in SI). Comparison with the data provides a lower bound of 5 accessible binding sites per vimentin filament cross-section (∼40 monomers (54)). While the microscopic interaction remains unresolved our data provide strong evidence for a direct and specific interaction between actin monomers and vimentin.

**Figure 5:**
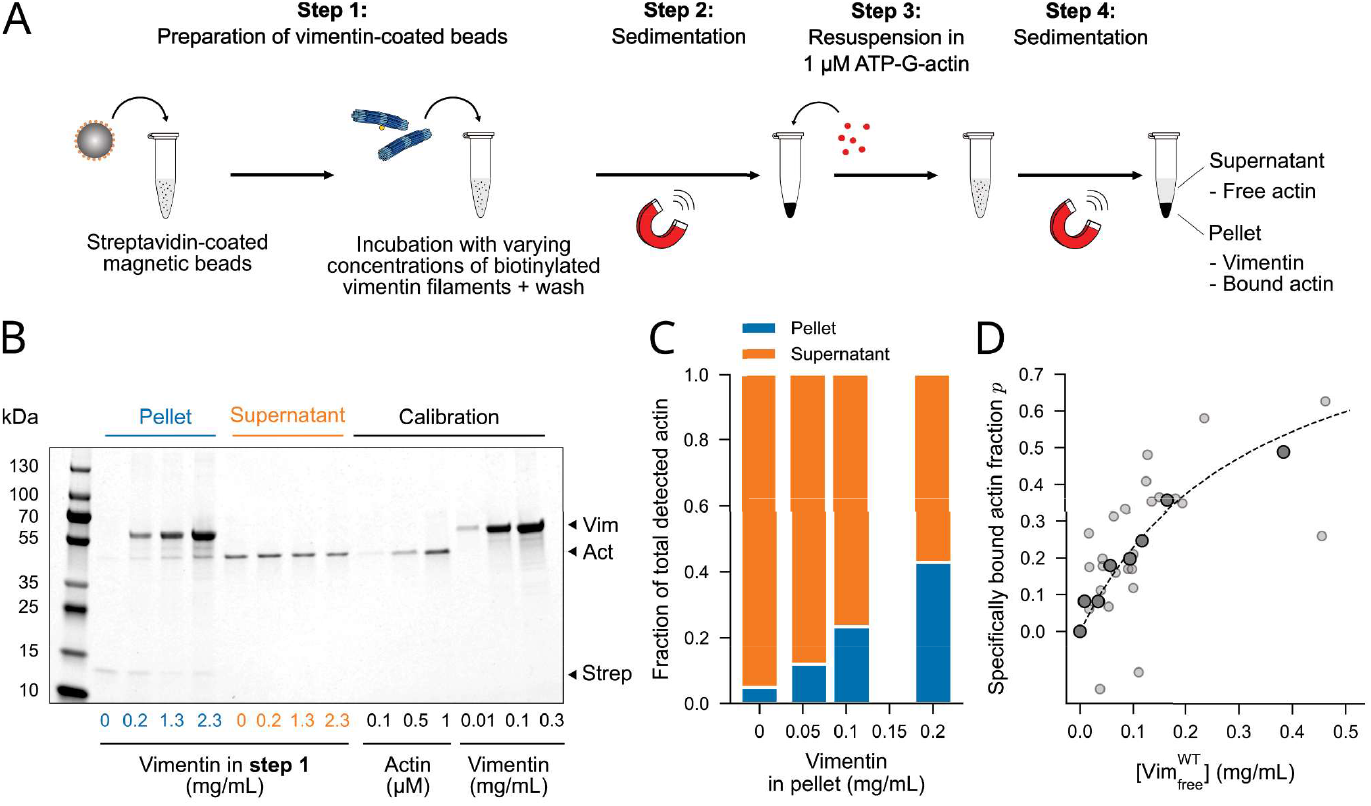
Vimentin directly interacts with actin. A. Actin-vimentin magnetic pull-down assay. In step 1, streptavidin-coated magnetic beads are incubated with varying concentrations of biotinylated vimentin filaments. In step 2, the beads are pulled with a magnet, and the supernatant containing unbound vimentin is discarded. The beads are further rinsed with buffer to completely remove unattached vimentin. In step 3, the vimentin-coated beads are resuspended in a solution containing 1 μM ATP-G-actin, and immediately pulled down with the magnet (step 4). The protein content of the pellet and the supernatant is then analyzed by SOS-PAGE. Step 3 and 4 were performed in less than a minute, limiting possible actin nucleation. The experiment was performed in the standard assembly buffer. B. Representative SOS-PAGE gel obtained with the assay described in A. Four concentrations of vimentin ranging from 0 to 2.3 mg/ml were used in step 1, and for each concentration, the pellet and supernatant obtained after step 4 were loaded. The six lanes on the right were loaded with samples containing known concentrations of actin or vimentin for calibration. Arrowheads indicate bands corresponding to vimentin (Vim), actin (Act), and streptavidin (Strep). The streptavidin band intensity provides a readout of the magnetic bead concentration. C. Quantification of the electrophoresis gel shown in B. Fraction of total detected actin found in the pellet (blue) and in the supernatant (orange) as a function of the vimentin concentration measured in the pellet. D. Fraction of available actin in solution specifically pulled with vimentin *(p)*, as a function of the free vimentin concentration in the pellet 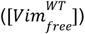. The amount of actin specifically bound to vimentin, as well as the amount of available actin in solution, was estimated by subtracting the amount of actin non-specifically bound to the beads, and by taking into account the difference in bead concentration between conditions (see Materials and methods in SI). The free vimentin concentration is defined as the total vimentin concentration measured in the pellet, minus the concentration of vimentin bound to actin. The latter is equal to the concentration of specifically bound actin, if we assume a 1:1 stoichiometry to extract the apparent G-actin-vimentin dissociation constant 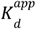 Data collected from N=S independent experiments were pooled together (small light dots) and binned by 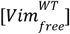 (large dark dots). Each bin contains four data points. The unbinned data were fitted (dashed line) to the equation: 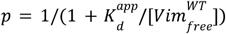.

## Discussion

In this work, we addressed the question of the direct interaction between actin and vimentin at the molecular level, providing new insight into cytoskeletal crosstalk. Combining in vitro assays with theoretical modeling, we found that the sole presence of vimentin enhances actin nucleation and barbed end elongation by specifically stabilizing ATP-actin subunits, without interfering with the binding of additional monomers. Pull-down experiments confirmed a direct interaction between the two proteins. Together, our findings identify vimentin as an unexpected regulator of actin dynamics and highlight a novel mechanism by which intermediate filaments can directly influence actin networks.

Our results rule out macromolecular crowding as an explanation of vimentin’s effect on actin dynamics. Previous in vitro studies using inert macromolecules have shown that macromolecular crowding can accelerate actin elongation, most likely due to excluded volume effects (55, 56). However, this effect requires concentrations of crowders typically above 20 mg/ml, a range we verified in our experimental conditions using Dextran (55, 56)(Fig. S7A). At 20 mg/ml, the volume occupied by Dextran macromolecules (40 kDa) can be estimated to be around 15%, assuming spheres with a hydrodynamic radius of 5 nm (57). In contrast, the accelerating effect of vimentin occurs at much lower concentrations (∼0.2 mg/ml) and quickly saturates at higher vimentin concentrations (∼1 mg/ml). In this range, the volume excluded by vimentin remains negligible (<0.5%) and cannot account for the observed acceleration. Moreover, the mechanism we uncovered - stabilization of ATP-actin at the barbed end through a decrease in the off-rate while leaving the on-rate unaffected - is fundamentally different from the effect of inert crowders. Indeed, the acceleration of actin elongation by these macromolecular crowders is dominated by an increase in the on-rate, which can be interpreted as an increase in the effective monomer concentration (56)(Fig. S7B). Finally, if excluded volume effects had a significant contribution to vimentin’s effect, one would expect vimentin’s effect to be independent of actin nucleotide state, and to observe the same acceleration for ADP-actin polymerization, which we did not observe. Altogether, these three observations - the low vimentin concentrations at which its effect occurs, the demonstrated stabilizing mechanism, and the dependency on actin nucleotide state - argue strongly against a role of crowding.

The specific stabilization of ATP-actin but not ADP-actin subunits at filament barbed ends by vimentin is an intriguing result. One possible explanation for this nucleotide dependence could be a difference in the accessibility of actin residues which can interact with vimentin. So far, only the structure of the barbed end with a terminal actin subunit in the ADP-state, which is in an F-actin conformation, has been obtained (58, 59). One can speculate that, during elongation, the recently added terminal subunit in the ATP-state could transiently remain in a G-actin-like conformation, which could thus allow it to interact with vimentin as supported by our pull-down assay. A second possible explanation for this nucleotide dependence is suggested by our observation that vimentin changes the critical concentration for actin elongation, and what this observation implies thermodynamically. Indeed, thermodynamics forbids the mere equilibrium binding of a protein with the filament barbed end from modifying this concentration, even though it can dramatically affect the individual on- and off-rate constants as long as their ratio remains unchanged. Thus, the change in the critical concentration induced by vimentin implies that vimentin’s effect operates out of equilibrium, which could be explained by a coupling of vimentin binding to the nucleotide dynamics, among other possible mechanisms.

The domain(s) of vimentin implicated in its effect on actin elongation remain unknown, although we ruled out the implication of vimentin head and tail domains in the experiments using truncated constructs. Mapping the potential interaction sites, for example, by systematic point mutation within the rod domain, crosslinking, or using structural approaches, would be critical to understand the actin-vimentin binding interface and how vimentin specifically stabilizes ATP-actin subunits at the barbed end. Cryogenic electron microscopy has been instrumental in understanding the mechanisms involved in actin assembly (40, 59, 60) and how it is regulated by actin-binding proteins at the barbed end (58, 61-63). Similar high-resolution structural approaches applied to F-actin and vimentin are required to confirm if vimentin filaments interact with the barbed end, and if so, if they act by creating additional bonds between actin subunits at the barbed end or by modulating their conformation.

We quantified two dissociation constants: one characterizing the vimentin/ATP-G-actin interaction (Fig. 5D) and one characterizing a putative interaction between vimentin ULFs and actin barbed ends (Fig.3D). However, it is difficult to compare these two dissociation constants quantitatively, as we do not resolve the involved microscopic interactions, which might differ between the two cases. Notably, the actin subunits at the barbed end and G-actin monomers expose distinct surface residues. Furthermore, the putative interaction between barbed ends and vimentin ULFs may involve multiple binding sites, possibly with some cooperativity.

Interestingly, recent in vitro studies demonstrated that vimentin stabilizes dynamic microtubules (64). The authors related this effect to a weak and transient microtubule-vimentin interaction, which they measured using optical tweezers. This interaction is sufficient to have a stabilization effect on microtubule dynamics without inducing co-alignment or bundling between the two filament types. Similarly, we propose that frequent but transient interactions of vimentin in the vicinity of the barbed end of actin filaments underlie the stabilizing effect that we have characterized. In contrast to its effect on actin, vimentin does not increase the microtubule elongation rate, but instead decreases the catastrophe frequency and increases the rescue frequency. The differences between vimentin’s effect on microtubules and on actin may arise from their distinct architectures and assembly pathways, although the underlying molecular mechanisms could be similar. However, because the study of the dynamic instability requires microtubules assembled from GTP-tubulin, the dependence of vimentin’s effect on the tubulin nucleotide state could not be directly assessed. Together, these findings suggest that vimentin may act as a broad modulator of cytoskeletal turnover.

Vimentin differs from canonical actin barbed end regulators such as formins and EnaNASP. These actin-binding proteins promote actin elongation through enhanced recruitment of (profilin-)actin monomers, thus having a major impact on the apparent actin on-rate (65–67). In contrast, vimentin does not significantly increase the recruitment of monomers but instead stabilizes the barbed end in the range of actin concentrations we tested. This suggests that vimentin does not act as a dominant driver of actin polymerization, such as formin and EnaNASP, but may rather provide a layer of fine-tuning that contributes to cytoskeletal remodeling in regions where vimentin, actin filaments, and regulatory factors converge. In cells, where the concentration of actin monomers is typically at least one order of magnitude higher than in our in vitro assay, the direct contribution of the off-rate reduction to actin elongation velocities may therefore be limited. However, the fact that vimentin directly impacts the barbed end raises the possibility that it could modulate the activity of actin-binding proteins by acting synergistically or antagonistically with canonical barbed-end regulators. Future work should test whether vimentin synergizes or competes with other actin regulators and how cross-linkers or signaling pathways could further tune this effect in cells.

## Materials and Methods

Detailed protocols are included in the SI Appendix. We purified recombinant wild-type human vimentin and vimentin mutants from E. coli (8, 9, 37). Vimentin, stored in 8M urea at -70 °C, was renatured by stepwise dialysis to sodium phosphate buffer (pH 7.0, 2.5mM sodium phosphate, 1 mM DTT) then prepared in a buffer compatible with actin assembly (assembly buffer: 10 mM HEPES pH 7.0, 1 mM sodium phosphate, 1 mM MgCl_2_, 0.2 mM EGTA, 100 mM KCI). KCI was added at the last minute to start vimentin filament assembly. The vimentin mix, typically at 1 mg/ml, was then incubated at 37°C for 30 minutes. We followed the same steps for experiments with vimentin Y117L and vimentin Δhead. The assembly protocol had to be optimized for vimentin Δtail to prevent the formation of unwanted aggregates. Vimentin Δtail filaments were assembled in a modified assembly buffer (10 mM HEPES, pH 7.0, 1 mM sodium phosphate, 1 mM DTT, 0.2 mM EGTA, 50 mM KCI), supplemented with 1 mM MgCl_2_ just before introducing the sample into the flow chamber. For ADP-actin polymerization experiments in Tris buffer, vimentin dialysed in Tris dialysis buffer (5 mM Tris-HCI pH 8.0, 1 mM DTT) was prepared in Tris assembly buffer (10 mM Tris-HCI, pH 7.0, 1 mM MgCl_2_, 0.2 mM EGTA, 50 mM KCI). The vimentin mix was then incubated at 37°C for 1 h.

Alpha skeletal actin was purified from rabbit muscle and stored in G-buffer (2 mM Tris-HCI, pH 7.8, 1 mM OTT, 0.2 mM CaCl_2_, 0.2 mM ATP, 0.01% NaN_3_)(37, 39). Actin was labeled with Alexa Fluor (AF)-488, Atto-643 or Atto-488 NHS Ester.

Polymerization and depolymerization experiments in TIRF microscopy were performed in flow chambers made of two cleaned glass coverslips (22 × 40 mm, #1.5, VWR) silanized with dichlorodimethylsilane. The flow chamber was first coated with neutrAvidin at 10 µg/ml for 5 minutes and rinsed with the assembly buffer. The surfaces were then passivated by incubation of Pluronic F127 1% for 15 min, thoroughly rinsed with the assembly buffer, and further passivated with bovine serum albumin (BSA) at 10 mg/ml for 5 min. Both F127 and BSA solutions were diluted in the assembly buffer. Biotinylated F-actin fragments were prepared by pre-assembly of F-actin (10% Atto-643-labeled, 10% biotinylated) in a test tube at 10 µM for 1 hour at room temperature (in G-buffer supplemented with 100 mM KCI), followed by a 1:500 dilution in assembly buffer containing Atto-643-labeled ATP-G-actin (non-biotinylated) at 0.2 µM to prevent depolymerization. The actin filaments were finally fragmented by strong pipetting and introduced into the flow chamber for a few seconds. The chamber was rinsed with the assembly buffer containing 0.2 µM Atto-643-labeled ATP-G-actin to remove unbound filaments. The polymerization experiment was then started by flowing 10 % AF 488 or Atto 488-labeled, 0.15 % biotinylated ATP-G-actin in the assembly buffer (except for AOP-G-actin polymerization) supplemented with 0.2 mM ATP, as well as 1 mM OTT, 1.2 mg/ml glucose, 40 µg/ml glucose oxidase, and 8 µg/ml catalase used as oxygen scavengers to limit photobleaching and creation of reactive oxygen species. The AOP-G-actin polymerization assay was performed in the Tris assembly buffer, supplemented with 1 mM OTT and the oxygen scavengers. Observations were carried out on an inverted microscope (Nikon Eclipse Ti), using TIRF (llas 2 system, Gataca systems) or wide field microscopy with a 60X objective NA 1.49. Images were acquired by a Kinetix camera (Teledyne Photometrics). The experiment was controlled using MicroManager. Movies were analyzed using TSOAX (68).

## Acknowledgements

We thank the Romet-Lemonne/Jegou lab for assistance and exciting discussions and specifically Ingrid Billaut-Chaumartin for careful reading of the manuscript. We thank Sarah Koster and Peter Bieling for sharing plasmids. CL thanks Harald Herrmann and Tatjana Wedig for teaching her vimentin purification, and the support of the EU-supported Cooperation in Science and Technology (COST) action NANONET. This project was funded by “lnvestissements d’Avenir” Labex WhoAml? (ANR-11-LABX-0071) and the Universite Paris Cite ldEx (ANR-IOEX-0001), the Agence Nationale pour la recherche (ANR-21-CE11-0004-02), the lmpulscience program of Fondation Bettencourt-Schueller and the Centre National de la Recherche Scientifique. LP was funded by La Ligue centre le Cancer (LNCC 282810).

## Supporting Information for

### Supporting figures

**Figure S1:**
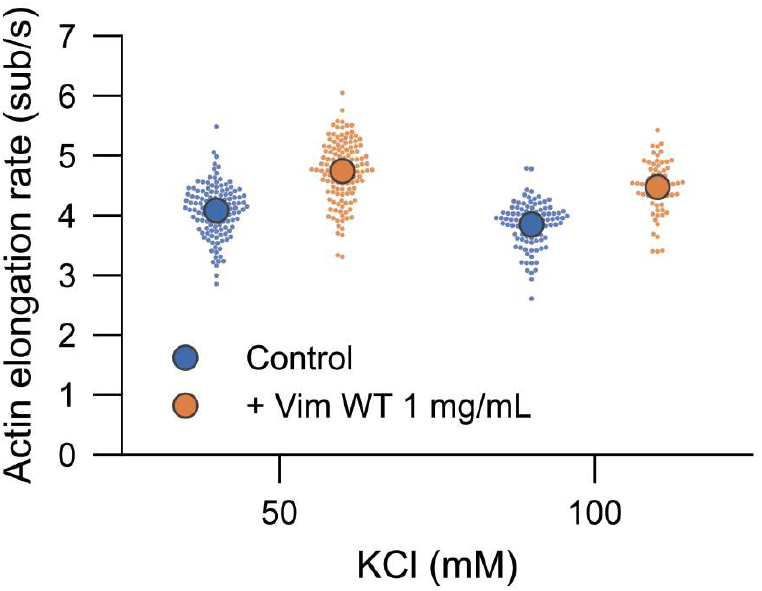
Vimentin has a similar impact on actin assembly at 50 and 100 mM KCI. Actin elongation rate in the absence (control, blue) or in the presence(+ Vim WT 1 mg/ml, orange) of vimentin filaments acquired in conditions using 50 mM and 100 mM KCI in the assembly buffer. Small dots: individual actin filament elongation rates (n ≥ 60 rates per condition), large dots: average elongation rates. The ATP-G-actin concentration is 0.5 µM.

**Figure S2:**
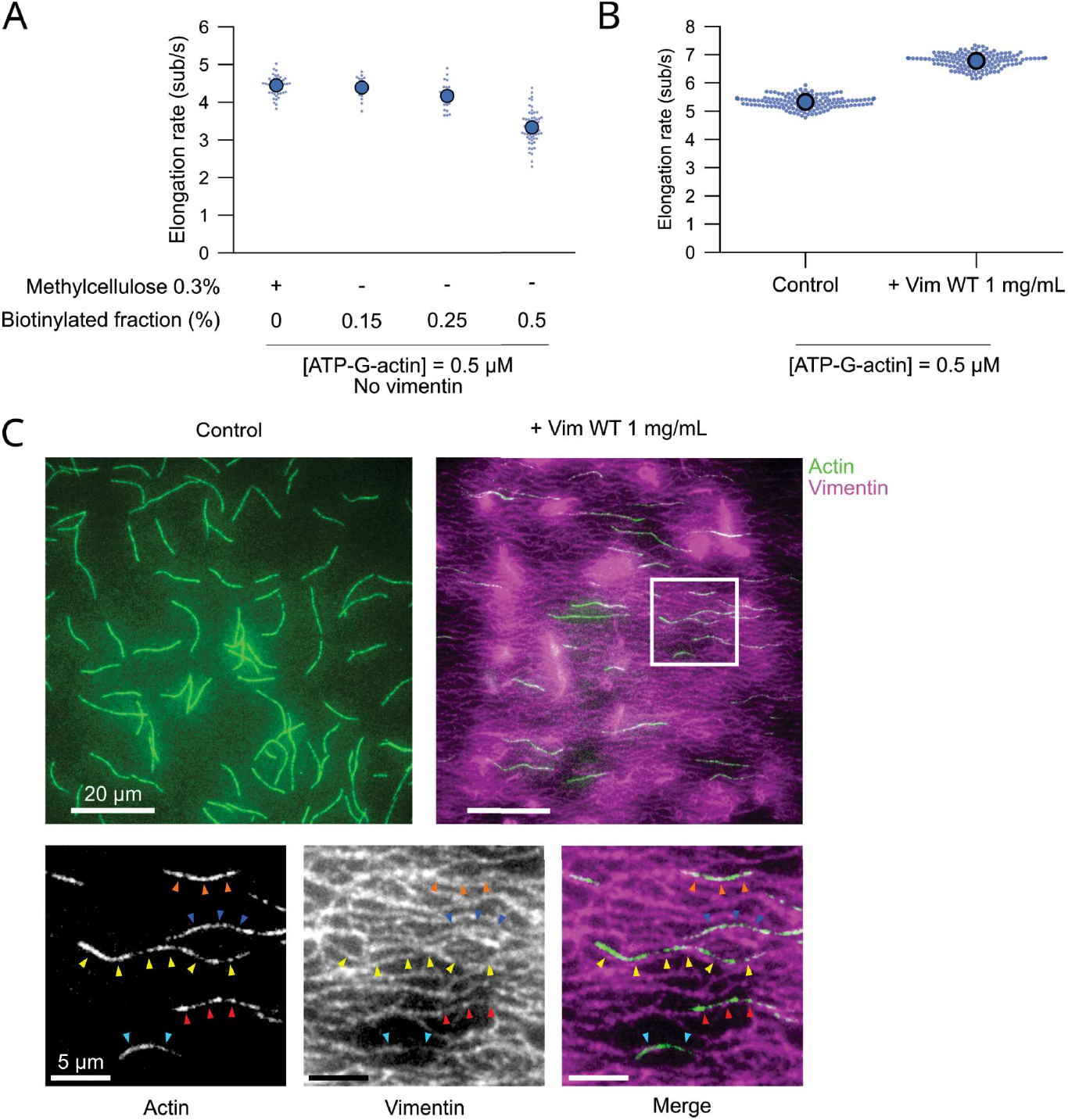
Impact of filament anchoring and methylcellulose. A. Impact of actin filament anchoring on the elongation rate. The elongation of actin filaments was monitored by TIRF microscopy, either keeping non-biotinylated filaments close to the surface using methylcellulose (0.3%) or using ATP-G-actin with varying biotinylation fractions and no methylcellulose. Small dots: individual actin filament elongation rates (n≥18 filaments), large dots: average elongation rates. B. Actin elongation rate in the absence (control) or in the presence(+ Vim WT 1 mg/ml) of pre-formed vimentin filaments in the methylcellulose conditions (0.3% methylcellulose, non-biotinylated actin filaments). Small dots: individual actin filament elongation rates (n≥140 filaments), large dots: average elongation rates. C. TIRF microscopy images of 10% labeled actin filaments (green) polymerizing in the presence of 0.3% methylcellulose, either in the absence (control) or in the presence (+ Vim WT 1 mg/ml) of pre-formed 2% labeled vimentin filaments (magenta). Enlarged images (bottom) show actin filaments aligned with the vimentin network, as emphasized by coloured arrow heads. The ATP-G-actin concentration in A, B and C is 0.5 µM.

**Figure S3:**
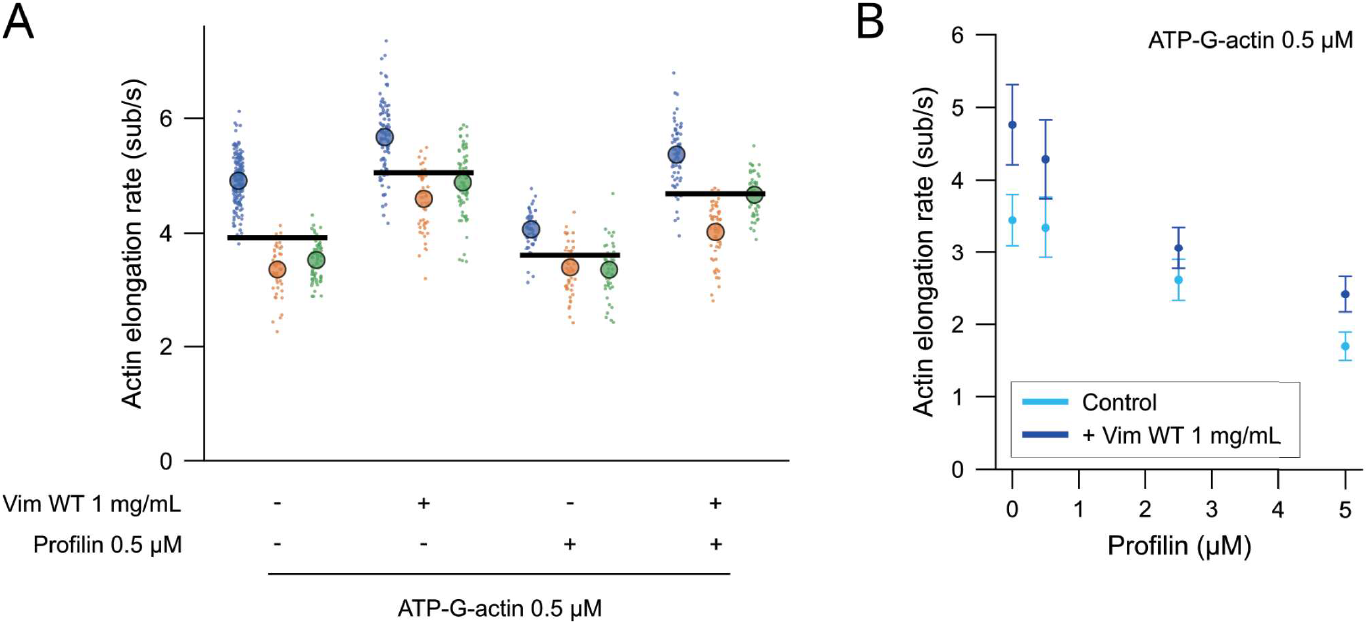
Vimentin promotes actin filament elongation from profilin-G-actin. A Actin elongation rate in the absence or presence of 1 mg/ml vimentin filaments (Vim WT), either with or without profilin 0.5 µM. The ATP-G-actin concentration was 0.5 µM. Small dots: individual actin filament elongation rates *(n ≥* 43 rates per condition and replicate), large dots: average elongation rates of the replicates *(N* = 3 independent replicates), black bars: mean elongation rate computed from the replicate averages. B. Actin elongation rate as a function of the profilin concentration, in the absence (light blue) or presence (dark blue) of 1 mg/ml vimentin filaments (Vim WT). The ATP-G-actin concentration was 0.5 µM. Data are presented as mean ± standard deviation (1 experiment, n ≥46 rates per condition).

**Figure S4:**
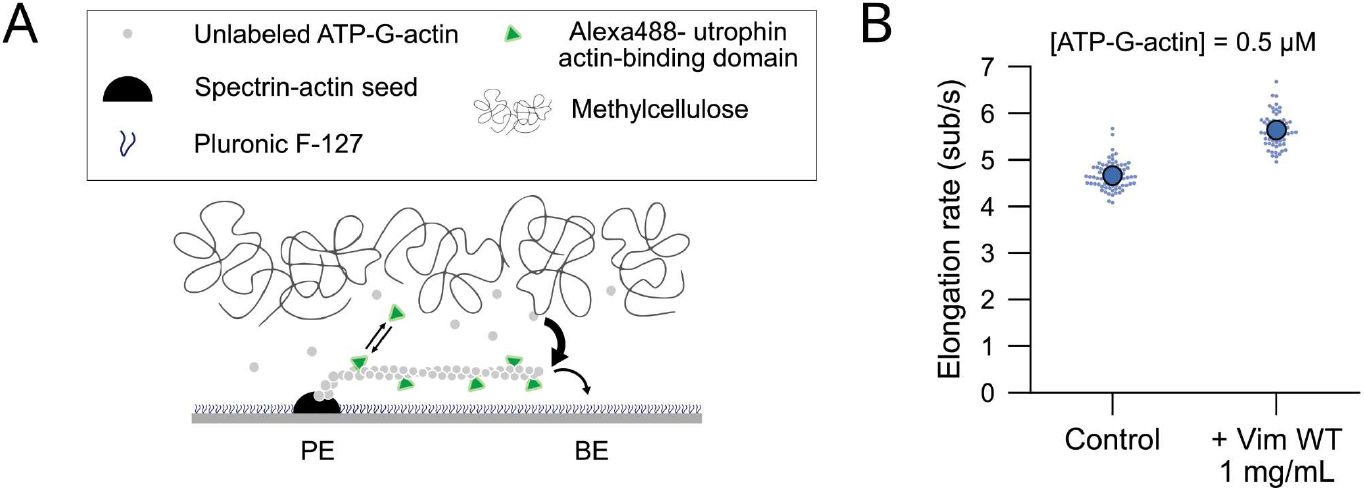
Utrophin does not interfere with the effect of vimentin on actin assembly. A. Actin assembly TIRF assay using utrophin actin-binding domain. The assembly of unlabeled ATP-G-actin (0.5 µM) from spectrin-actin seeds was monitored by TIRF microscopy using Alexa488-utrophin actin-binding domain (50 nM) to label the filaments. The filaments were maintained close to the surface using 0.3% methylcellulose. B. ATP-actin elongation rate in the absence (control) or in the presence(+ Vim WT 1 mg/ml) of pre-formed vimentin filaments in this utrophin-based TIRF assay. Small dots: individual actin filament elongation rates (n ≥ 63 filaments), large dots: average elongation rates.

**Figure S5:**
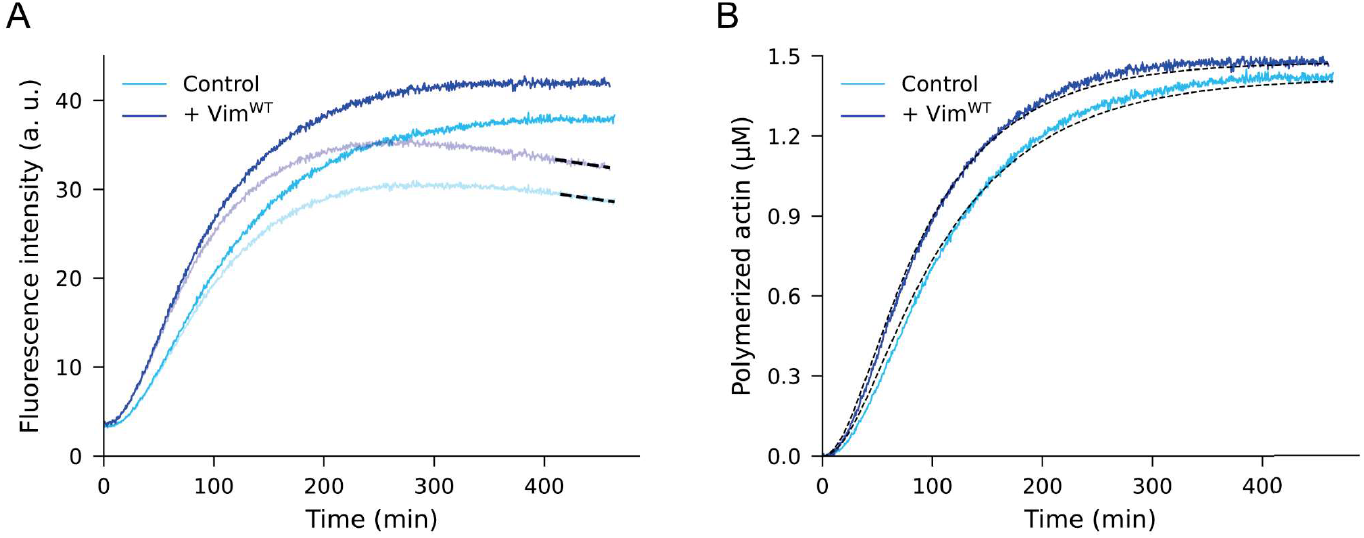
Quantification of the nucleation rate constant using our theoretical model. Illustration of the key steps used to theoretically predict the nucleation rate constant within our model, performed on the representative replicate of Fig. 4. Actin (1.5 µM, 8% pyrene-labeled) was assembled in the absence (Control, light blue) and in the presence of 1 mg/ml of pre-assembled wild-type vimentin filaments(+ Vim^WT^, dark blue). A. Correction of the fluorescence data for photobleaching. The fluorescence decrease at long times in the fluorescence data (transparent curves) was fitted with an exponential (dashed lines) to obtain the bleaching rate under Control and + Vim^WT^ conditions, respectively. The fluorescence curves were subsequently corrected for photobleaching based on Rosenbloom et al (1) to enable further analysis steps (plain curves, see Materials and methods). B. Fit of the theoretical model described in the main text to the polymerized actin concentration determined from the fluorescence data (also see Theoretical modeling). After correcting for photobleaching, the fluorescence curves from A were multiplied by a scaling factor to translate the fluorescence intensity into concentration of polymerized actin (plain curves, see Materials and methods). The polymerization curves were fitted with the model prediction for the polymerized actin concentration (Eq. (11), dashed lines) to determine the nucleation rate constant, which is the only free parameter.

**Figure S6:**
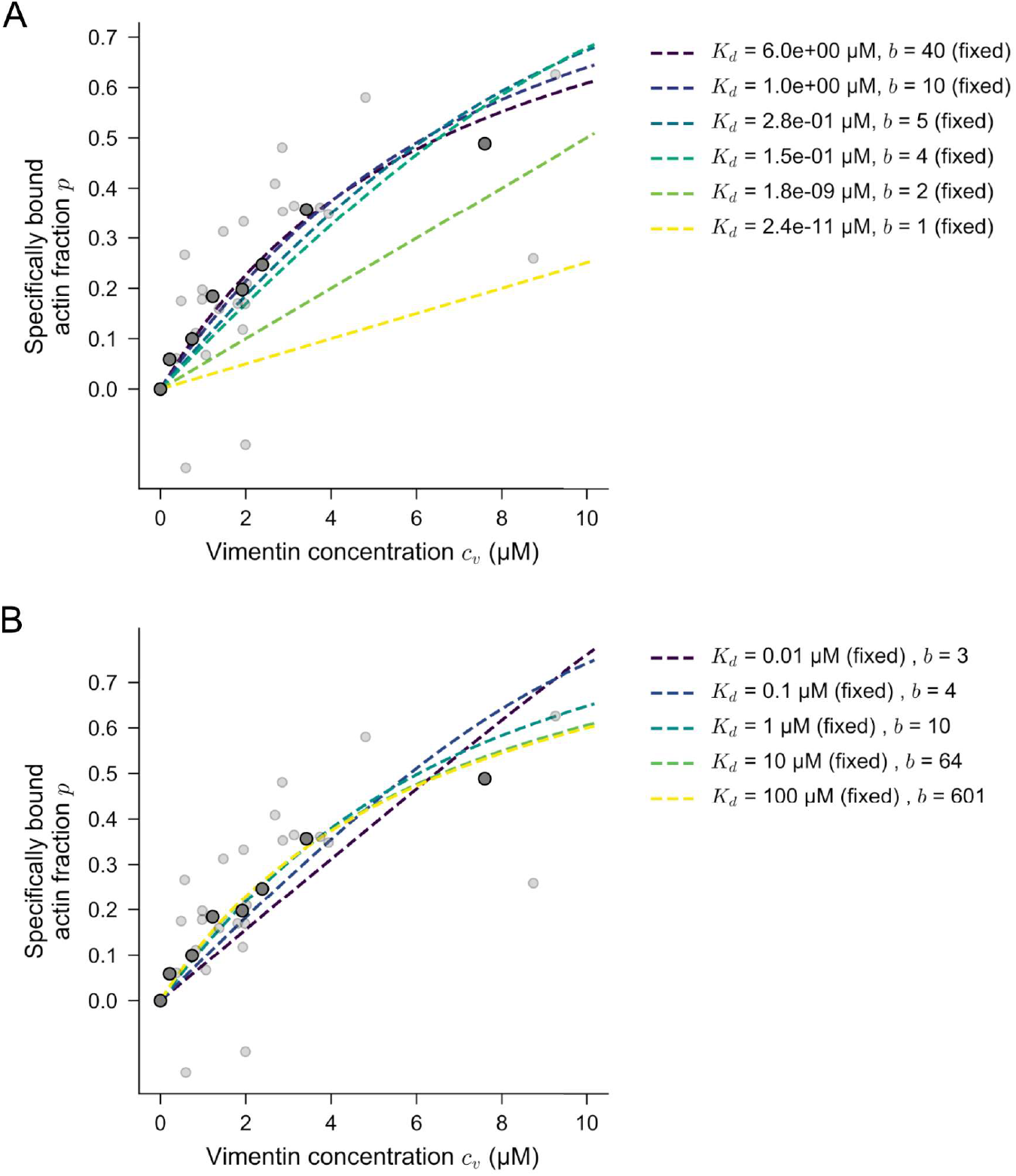
Each vimentin cross-section has multiple binding sites for G-actin. Fits of a thermodynamic binding model to the fraction of *p* actin available in solution specifically pulled down with vimentin, as a function of the vimentin concentration in the pellet *c_v_* expressed in µM of vimentin monomers. The total G-actin concentration is fixed to *c_a_* = 1 µM. Data collected from N=5 independent experiments were pooled together (small light dots) and binned by *c_v_* (large dark dots; each bin contains four data points). *b* is the number of accessible binding sites per vimentin cross-section. *K*_*d*_ expressed in µM of vimentin monomers, denotes the ATP-G-actin-vimentin dissociation constant, or equivalently the concentration of free vimentin binding sites needed to bind half of the actin monomers. A. Dashed lines: One-parameter fits of Eq. (20) to the unbinned data, using *K*_*d*_ as the only free parameter. For each fit, we fix *b* to a different value given in the legend. B. Dashed lines: One-parameter fits of Eq. (20) to the unbinned data, using *b* as the only free parameter. For each fit, we fix *K*_*d*_ to a different value given in the legend.

**Figure S7:**
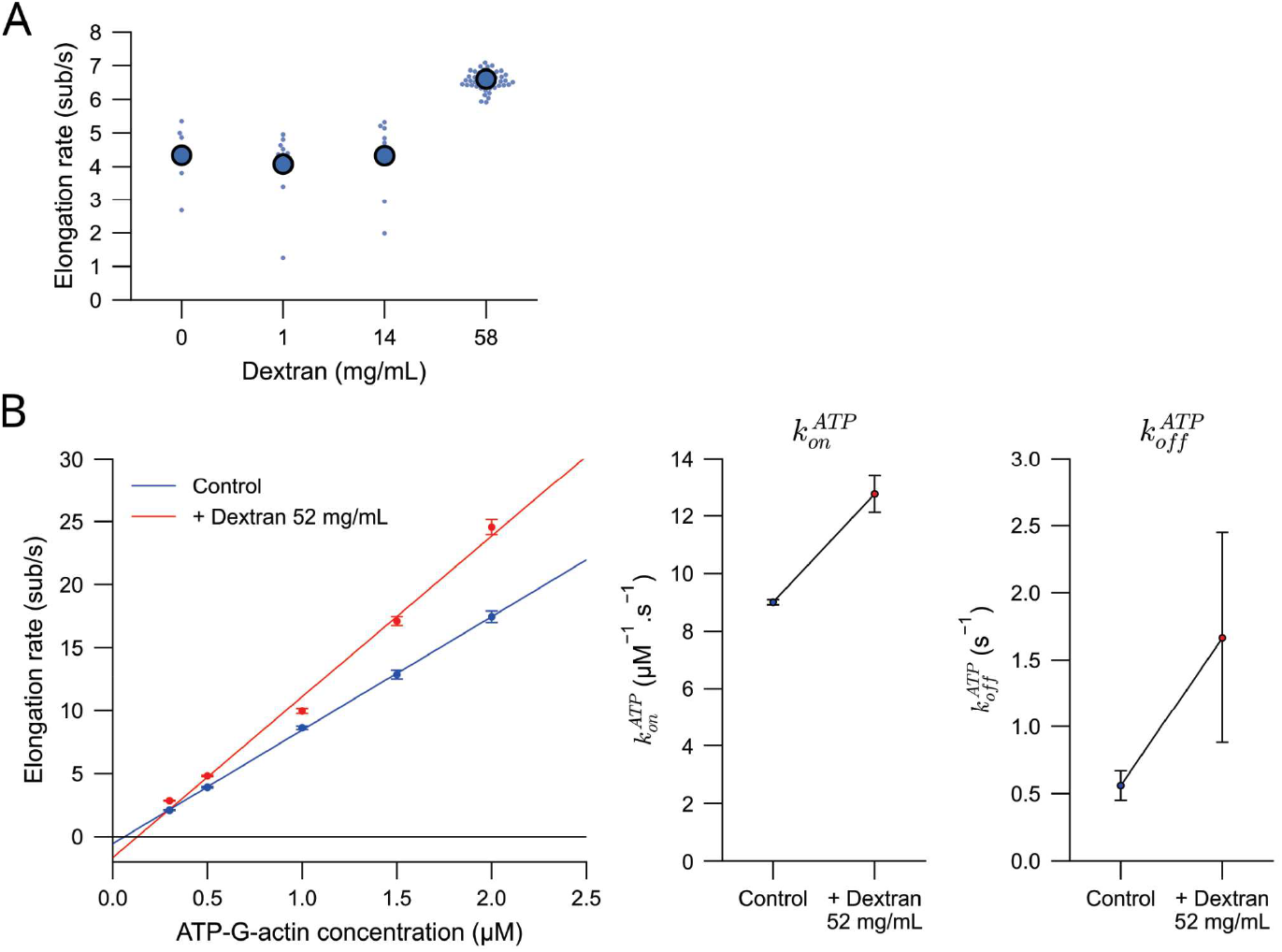
Effect of Dextran on actin assembly. A. Barbed end elongation rate in the presence of varying Dextran (40 kDa) concentrations. In this experiment, neither methylcellulose nor biotinylated actin were used, and only filaments growing parallel to the surface for a long enough time could be tracked. Note that in the presence of a high Dextran concentration (> 50 mg/ml), actin filaments were kept close to the surface just like when using methylcellulose, allowing to track more filaments. Small dots: individual actin filament elongation rates (n ≥ 7 filaments), large dots: average elongation rates. The ATP-G-actin concentration is 0.5 µM. B. Impact of a high Dextran concentration (52 mg/ml) on actin on- and off-rates. Left: actin elongation rate as a function of ATP-G-actin concentration in the absence (control, blue) or in the presence of Dextran (+ Dextran 52 mg/ml, red). Data are shown as the mean ± standard error of the mean (1 experiment, n ≥ 23 filaments per condition). Solid lines represent linear fits of the means (similar to Fig. 1E) to extract the on- and off-rates 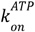 and 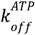 under both conditions. Right: measured actin on- and off-rates 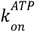 and 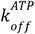 in the absence (control, blue) or in the presence of Dextran (+ Dextran 52 mg/ml, red). Error bars represent the square root of the variances of the estimated parameters, extracted from the covariance matrix returned by the numpy.polyfit function in Python. 0.15%-biotinylated ATP-G-actin was used in the control condition.

**Figure S8:**
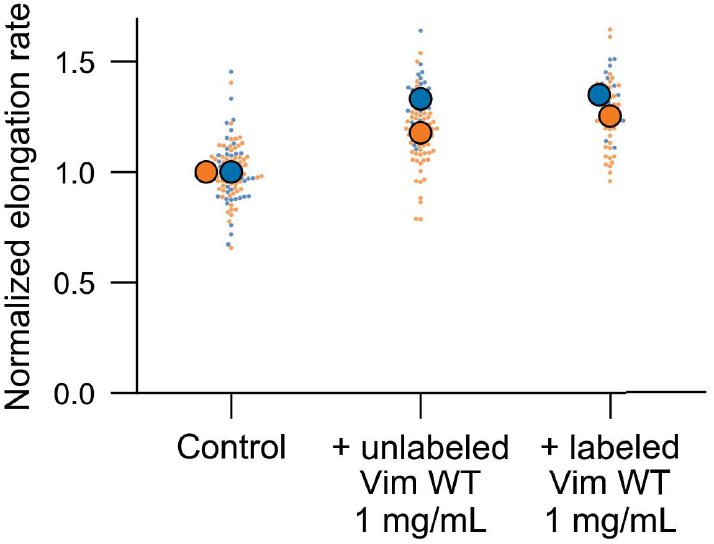
Vimentin labeling has no impact on its effect on actin assembly. Normalized actin elongation rate in the absence (control) or in the presence of unlabeled or 2% labeled pre-assembled vimentin filaments (+ (un)labeled Vim WT 1 mg/ml). Colors represent experiments performed on the same day. The rates were normalized with respect to the average rate measured in the control condition on the same day. Small dots: individual actin filament elongation rates (n ≥ 13 filaments), large dots: average elongation rates. The ATP-G-actin concentration is 0.5 µM.

**Figure S9:**
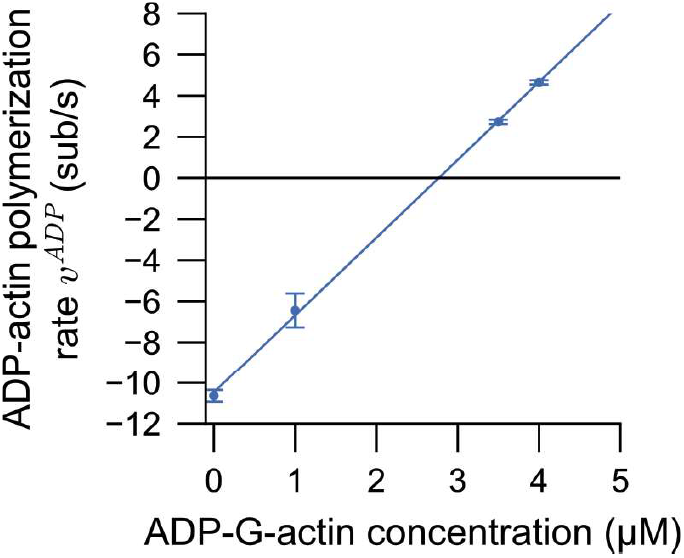
ADP-actin polymerisation in Tris buffer. Control (de) polymerization experiments in phosphate-free buffer were performed in Tris assembly buffer (10 mM Tris-HCI, 1 mM MgCl_2_, 0.2 mM EGTA, 50 mM KCI). ADP-actin polymerization rate as a function of the ADP-actin concentration. Data are shown as the mean ± standard error of the mean (n ≥ 4 filaments per condition). Solid line represents linear fit of the means (similar to Fig. 1E) to extract the on- and off-rates 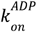 and 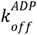 in these phosphate-free conditions. The estimated on- and off-rates are 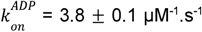 and 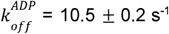, respectively (estimated parameter ± square root of the variance of the estimated parameter, extracted from the covariance matrix returned by the numpy.polyfit function in Python).

## Materials and Methods

### Vimentin constructs

The full-length human vimentin cDNA subcloned into the prokaryotic expression vector pDS5 with bacterial expression was a kind gift from Harald Herrmann (2). Vimentin Δtail and vimentin Δhead constructs were produced from the full-length vimentin plasmid using the Q5 Site-Directed Mutagenesis Kit (catalog no.E0554S, Biolabs) using the following primers: Δtail, forward primer TAAAAATTGCACACACTCAGTGC and reverse primer AATCCTGCTCTCCTCGCC (removal of last 55 amino acids); Δhead forward primer CGGCTCCTGCAGGAC and reverse primer CGGCTCCTGCAGGAC (removal of the first 76 amino acids after the methionin). Vimentin-Y117L plasmid for bacterial expression was kindly provided by Sarah Koster (3).

### Vimentin purification (WT, Y117L, Δtail, Δhead) and labeling

Vimentin WT and mutants were purified from E. coli bacteria in denaturating conditions, as previously described (4-7) and stored in a buffer containing 8 M urea, 5 mM TRIS, 1 mM EDTA, 0.1 mM EGTA, and 10 mM methylamine hydrochloride solution, pH 7.5 at -70°C.

Before experiments, vimentin stock solution in urea was transferred to a dialysis tubing (Servapor, cutoff at 12 kDa) and renatured by stepwise dialysis from 8, 6, 4, 2, 1, 0 M urea into sodium phosphate buffer (2.5 mM, pH 7.0, 1 mM OTT) with 15 min for each step. The final dialysis step was performed overnight at 4 °C with 2 L of the sodium phosphate buffer. Renatured proteins were stored at 4°C for a maximum of 10 days. The vimentin concentration was determined by UV-VIS absorbance measurements using a molecular weight of 53.65 kDa and an extinction coefficient at 280 nm of 22,450 M-^1^cm-^1^ (8). After dialysis, vimentin was either directly used for experiments or flash frozen in liquid nitrogen and stored at -70°C.

We labeled vimentin proteins by forming a maleimide-thiol covalent bond between an Alexa Fluor 555 or 568 (C2 maleimide, Thermofisher) or Maleimide-PEG2-Biotin EZ-Link (Thermofisher) and the cysteine-328 on vimentin as previously described (4-6, 9). We used a labeling fraction of 2% to get an idea of the vimentin network assembly and verified that labeled and unlabeled networks gave a similar effect on actin elongation (Fig. S8). We systematically used a 2% labeling fraction of vimentin Δtail to be sure that the network assembled correctly without forming aggregates. In these experiments, the effect of 2%-labeled vimentin Δtail on actin assembly was compared to the effect of 2%-labeled WT vimentin. Depolymerization experiments in Fig. 2A-B were performed using 2-5%-labeled WT vimentin.

### Vimentin assembly

For the actin polymerization/depolymerization TIRF assays, vimentin was first renatured in 2.5 mM sodium phosphate (pH 7) supplemented with 1 mM OTT and then prepared in a buffer compatible with actin assembly (assembly buffer: 10 mM HEPES pH 7.0, 1 mM sodium phosphate, 1 mM MgCl_2_, 0.2 mM EGTA, 100 mM KCI). KCI was added at the last minute to start the assembly, using the so-called kick-start method. The vimentin mix, typically at 1 mg/ml, was then incubated at 37°C for 30 minutes. We followed the same steps for experiments with vimentin Y117L and vimentin Δhead. For ADP-actin polymerization experiments in Tris buffer, vimentin dialysed in Tris dialysis buffer (5 mM Tris-HCI pH 8.0, 1 mM OTT) was prepared in Tris assembly buffer (10 mM Tris-HCI, pH 7.0, 1 mM MgCl_2_, 0.2 mM EGTA, 50 mM KCI). The vimentin mix was then incubated at 37°C for 1 h.

The assembly protocol had to be optimized for vimentin Δtail to prevent the formation of unexpected aggregates. Vimentin Δtail was ultracentrifuged for 20 min at 4°C and 56,000 rpm prior to experiments. Filaments were then assembled in a modified assembly buffer (10 mM HEPES, pH 7.0, 1 mM sodium phosphate, 1 mM OTT, 0.2 mM EGTA, 50 mM KCI). The buffer was supplemented with 1 mM MgCl_2_ just before introducing the sample into the flow chamber. The KCI concentration was kept at 50 mM for these experiments and corresponding assays in the control condition and with vimentin wild-type.

For the pull-down assay, we assembled vimentin filaments by the addition of KCI to a final concentration of 100 mM to a solution of freshly dialyzed vimentin tetramers in 2.5 mM sodium phosphate, pH 7.

### Actin purification and labeling

Alpha skeletal actin was purified from rabbit muscle as previously described (10) and stored in G-buffer (2 mM Tris-HCI, pH 7.8, 1 mM DTT, 0.2 mM CaCl_2_, 0.2 mM ATP, 0.01% NaN_3_)(6, 11). Actin was labeled with Alexa Fluor (AF)-488, Atto-643 or Atto-488 NHS Ester (Thermofisher) following provider’s instructions. Spectrin-actin seeds were purified from human erythrocytes (12).

### Preparation of ADP-actin

ADP-actin was obtained by incubating ATP-actin at 10 µM in ATP-free G-buffer supplemented with 2.5 mM glucose, 0.2 mM EGTA, 20 µM MgCl_2_, and 15 U/ml hexokinase for 18 h on ice. This protocol was adapted from (13, 14). ADP-actin was then used for experiments within a few hours. Complete conversion of ATP-actin into ADP-actin was verified by determining the on- and off-rate at the barbed end and which were consistent with those reported in the literature in similar buffer conditions (15)(Fig. S9)

### Actin polymerization and depolymerization TIRF assays

Polymerization and depolymerization experiments in TIRF microscopy were performed in flow chambers made of two cleaned glass coverslips (22 × 40 mm, #1.5, VWR) silanized with dichlorodimethylsilane following previously published detailed protocols and separated by melted parafilm strips (16). In short, we first cleaned coverslips by washing and sonicating them in a series of Hellmanex, KOH, and ethanol solutions. The coverslips were then completely dried and treated in a plasma cleaner for 5 min. Immediately after plasma treatment, the coverslips were soaked in dichlorodimethylsilane 0.05% in trichloroethylene and incubated at room temperature for 1 h. We then washed off the excess silane by sonicating the coverslips 3 times in methanol for 15 min each. The silanized coverslips were dried by compressed air and stored at 4 °C for no longer than 2 weeks. This silanization method creates strongly hydrophobic surfaces, which we use to strongly adsorb neutravidin in combination with blocking the rest of the surface with Pluronic F127. The resulting flow chambers between the parafilm strips had a width of about 2 mm.

The flow chamber was first coated with neutrAvidin at 10 µg/ml for 5 minutes and rinsed with assembly buffer (10 mM HEPES, pH 7.0, 1 mM sodium phosphate, 1 mM MgCl_2_, 0.2 mM EGTA, 100 mM KCI). The surfaces were then passivated by incubation of Pluronic F127 1% for 15 min, thoroughly rinsed with the assembly buffer, and further passivated with bovine serum albumin (BSA) at 10 mg/ml for 5 min. Both F127 and BSA solutions were diluted in the assembly buffer. Biotinylated F-actin fragments were prepared by pre-assembly of F-actin (10% Atto-643-labeled, 10% biotinylated) in a test tube at 10 µM for 1 hour at room temperature, followed by a 1:500 dilution in assembly buffer containing Atto-643-labeled ATP-G-actin (non-biotinylated) at 0.2 µM to prevent depolymerization. The actin filaments were finally fragmented by strong pipetting and introduced into the flow chamber for a few seconds. The chamber was rinsed with assembly buffer containing 0.2 µM Atto-643-labeled ATP-G-actin to remove unbound filaments. The polymerization experiment was then started by flowing 10 % AF 488 or Atto 488-labeled, 0.15 % biotinylated ATP-G-actin in assembly buffer supplemented with 0.2 mM ATP, as well as 1 mM DTT, 1.2 mg/ml glucose, 40 µg/ml glucose oxidase, and 8 µg/ml catalase used as oxygen scavengers to limit photobleaching and creation of reactive oxygen species.

For polymerization and depolymerization of actin filaments attached to the surface by their pointed end, the silanized coverslips were first coated with spectrin-actin seeds, then passivated by F127 1% and BSA 10 mg/ml diluted in assembly buffer. Actin filaments were polymerized by flowing non-biotinylated ATP-G-actin in assembly buffer supplemented with 0.3% methylcellulose, 0.2 mM ATP, 1 mM DTT, 1.2 mg/ml glucose, 40 µg/ml glucose oxidase, and 8 µg/ml catalase. The depolymerization of actin filaments was monitored after rinsing the flow chamber with assembly buffer and assembly buffer supplemented with 0.3% methylcellulose, 0.2 mM ATP, 1 mM DTT, 1.2 mg/ml glucose, 40 µg/ml glucose oxidase, and 8 µg/ml catalase.

The depolymerization experiment in Fig. 2A was performed in our standard assembly buffer, which contains 1 mM sodium phosphate. This phosphate concentration is close to the affinity constant of inorganic phosphate for actin within the filament (1.5 mM, (15, 17)), meaning that a substantial amount of actin subunits in the filament are in the ADP.P; state. Consistently, the barbed end depolymerization rate measured in the control condition was 3.4 ± 0.7 sub/s (mean ± SD), which lies between the values expected for ADP-actin (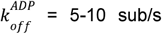, (14, 15, 11), (Fig. S9)) and ADP.P;-actin (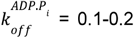 sub/s (11, 15). We thus measured an effective depolymerization rate resulting from the alternative unbinding of ADP and ADP.P, subunits, which we denoted 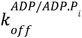.

The depolymerization experiment in Fig. 2B was performed in the Tris assembly buffer supplemented with 1 mM DTT, 1.2 mg/ml glucose, 40 µg/ml glucose oxidase, and 8 µg/ml catalase.

ADP-actin polymerization experiments (Fig. 2C) were performed in the Tris assembly buffer supplemented with 1 mM DTT, 1.2 mg/ml glucose, 40 µg/ml glucose oxidase, and 8 µg/ml catalase. The polymerization of unlabeled ADP-actin was monitored by adding 50 nM Alexa488-labeled actin-binding domain of utrophin, following previous work (18, 19).

Observations were carried out at 25°C on an inverted microscope (Nikon Eclipse Ti), using TIRF or wide field microscopy with a 60X or 100x objective NA 1.49. Images were acquired by a Kinetix camera (Teledyne Photometrics). The experiment was controlled using MicroManager.

### Actin polymerization and depolymerization rate quantification

Polymerization and depolymerization movies were analyzed using TSOAX (20). Before automatic filament detection and tracking in TSOAX, the images were pre-processed on Fiji: the contrast was enhanced by using the Subtract Background function (rolling ball radius 50 pixels), and the drift was corrected using the Template Matching plugin. Output tracking data returned by TSOAX were then processed using a-homemade Python codes to extract actin assembly and disassembly rates.

To quantify the polymerization rates at the barbed end, we only considered filaments with a fixed end (either attached to a spectrin-actin seed or an anchored F-actin fragment) to avoid filaments potentially polymerizing from both ends. We also filtered out tracks where the filament was detected for less than 10 frames. We then implemented an automatic filtering of the tracks to remove filament detection errors, resulting in longer or shorter filaments to get a more accurate estimation of the (de)polymerization rate. For each track, we performed a linear fit on the elongation data (length as a function of time). The coefficient of determination (R^2^) was computed. Through an iterative process, we identified data points resulting from detection errors by, at each iteration, identifying which data point was yielding the highest R^2^ when removed. This iterative process was stopped as soon as one of the three following conditions was met: a) R^2^ ≥ 0.999, or b) the track had 10 data points left, or c) 30% of the data points of the track were removed. After this iterative process, the R^2^ was checked, and only tracks with R^2^ ≥ 0.99 were kept. Thus, only tracks where the filament was correctly detected for at least 10 frames were kept, and tracks where more than 30% of the points had to be removed to obtain a satisfactory fit were rejected.

Depolymerization experiments exhibited frequent pauses, likely due to photo-induced dimerization, and required a slightly different analysis routine. First, tracks where the filament was detected for less than 10 frames were removed. Then, we performed a linear fit on the elongation data (length as a function of time), computed the coefficient of determination (R2), and tracks with R^2^<0.7 were discarded. We then implemented a semi-automatic processing of the remaining tracks, based on the piecewise-regression Python package (21). For each track, the user decided whether to keep or discard it. For selected tracks, the user specified the number of breakpoints, and the piecewise-regression package was called to automatically segment the track in linear segments (pauses or depolymerization periods). The user then confirmed or rejected the segmentation. If confirmed, the slope was computed on each segment of the track. For each tracked filament, only segments longer than four time frames were considered, and the highest slope was selected as the depolymerization rate for this filament.

For each experiment, condition, and replicate, outliers among measured rates were filtered out based on their Z-score (outlier if ∣Z∣-score > 3), and only replicates with at least 20 remaining measured rates after filtering were kept.

### Direct visualization of actin nucleation by fluorescence microscopy

For actin nucleation assays, glass coverslips (22 × 40 mm and 18 × 18 mm, #1.5, VWR) were first cleaned following the same protocol as for TIRF assays. We then pipetted drops of a Pll-PEG solution (0.1 mg/ml in 10 mM HEPES pH 7.4) and placed a cleaned glass coverslip on top of each drop to incubate for 1-2h (100 µl drops for 18 × 18 mm coverslips, and 200 µl for 22 × 40 mm coverslips). The coverslips were then rinsed with milliQ water and gently dried with a kimwipe (Kimtech). 10% Atto-488-labeled ATP-G-actin was mixed to a final concentration of 300 nM in assembly buffer or assembly buffer containing pre-assembled wild-type vimentin filaments at 1 mg/ml, supplemented with 0.5% BSA, 0.2 mM ATP, 1 mM DTT, 1.2 mg/ml glucose, 40 µg/ml glucose oxidase, and 8 µg/ml catalase. We then deposited 0.7 µl of this mixture on a Pll-PEG-coated coverslip (22 × 40 mm) and put an 18 × 18 mm coverslip on top, resulting in a chamber with a height of ∼2 µm. The chamber was sealed using mineral oil (M8410, Sigma-Aldrich), and the sample was imaged at different time points by fluorescence microscopy. The number of filaments per field of view was then counted manually.

### Pyrene assay

Bulk actin nucleation and polymerization kinetics were followed by fluorimetry in a SAFAS Xenius fluorimeter at room temperature. Pyrenyl-actin was made by labelling actin with N(1-pyrene)-iodoacetamide (Thermo Fisher Scientific) (22). ATP-G-actin (8% pyrene-labeled) was mixed to a final concentration of 1.5 µM in assembly buffer or assembly buffer containing pre-assembled wild-type vimentin filaments at 1 mg/ml. Control samples containing assembly buffer alone or assembly buffer with pre-assembled vimentin filaments at 1 mg/ml were also prepared. The samples were transferred into quartz cuvettes, and their fluorescence intensities were recorded as a function of time, using excitation and emission wavelengths of 366 and 407 nm, respectively. Because vimentin has its own fluorescence signal at these wavelengths, the fluorescence signals of the vimentin-containing solutions were corrected by subtracting the signal of the vimentin solution in assembly buffer. Accordingly, the fluorescence signals of the control solutions (actin in assembly buffer alone) were corrected by subtracting the signal of the assembly buffer. We determined the photobleaching rate by fitting the last 100 time points (around 50 minutes) of the representative fluorescence data in Fig. 4A with an exponential, obtaining one value for the photobleaching rate under control and vimentin conditions each (Fig. S5A). We corrected all replicates for photobleaching using these values (1). In order to convert the corrected fluorescence signals into concentrations of polymerized actin, we followed the method proposed by Doolittle et al. (23) with minor changes: the minimum intensity *I_mm_*. at time zero was obtained by fitting a straight line to the 10 earliest time points, and we used the mean of the last 30 time points as the maximum intensity *I_max_*. To compute the critical concentration,-determined in our buffer conditions by TIRF microscopy (Fig. 1F): 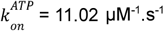 and 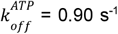 in the control condition, and 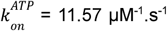 and 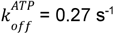 with vimentin. We extracted the empirical initial nucleation rate by adapting the steps in Doolitlle et al. (23), fitting a pure quadratic *At*^2^ to the concentration of polymerized actin to avoid overfitting, with a concentration cutoff y = 0.1.

### Pull-down assay

To probe the direct interaction between actin and vimentin, we implemented a magnetic pull-down assay. Streptavidin-coated magnetic beads (Dynabeads M-270, ThermoFisher) were thoroughly washed in the vimentin assembly buffer (2.5 mM sodium phosphate buffer, pH 7, provided with 100 mM KCI), and incubated with varying concentrations of biotinylated vimentin filaments (with ∼15% labeling fraction) assembled in the vimentin assembly buffer for 1 minute at 37°C. The vimentin-coated beads were then washed twice with 1 ml of vimentin assembly buffer to remove unattached vimentin. The vimentin beads were then pulled down with a magnet, and the supernatant was discarded. The beads were resuspended in a 1 µM ATP-G-actin solution in the assembly buffer and rapidly pulled down with a magnet. The pellet and the supernatant were separated within a few seconds with the magnet, and the pellet was resuspended for SOS-PAGE electrophoresis. We used 4-20% Mini-PROTEAN TGX precast polyacrylamide gels with 15 wells (Biorad). We mixed the samples with 6X Laemmli buffer provided with 10% beta-mercaptoethanol at a ratio of 5:1 and loaded 12 µL per well. We ran the gels for 60 min at 160 V with Tris/Glycine/SOS buffer (Biorad) and stained the gels overnight with lnstantBlue protein stain (Expedeon). The intensity of the bands observed on the SOS-PAGE gels was analyzed by plotting the intensity profiles of the lanes along lines of constant width using Fiji, and by fitting these intensity profiles with multiple Gaussians to extract the area under each peak. The concentrations of actin and vimentin were estimated by comparing the intensity of the bands with a range of actin and vimentin samples with known concentrations used for calibration. The amount of actin specifically bound to and pulled down with vimentin was estimated as follows. For each condition, the actin band intensity was divided by the bead band intensity to get the actin-per-bead intensity ratio. For each condition with vimentin, we subtracted the actin-per-bead intensity ratio of the control condition without vimentin, accounting for non-specific binding of actin on the beads. We then multiplied the result by the bead intensity to retrieve the amount of actin intensity corresponding to proteins that were specifically pulled down with vimentin. This actin intensity was then converted to an actin concentration using the calibration.

### Theoretical modeling

#### Actin nucleation and elongation with a limited monomer pool

Here we model F-actin nucleation and elongation subject to a limited pool of G-actin monomers. We obtain an analytical prediction of the time-dependent barbed end concentration (Fig. 4C) and the initial nucleation rate (Fig. 4B). To determine the only unknown parameter in our model, namely the nucleation rate constant, we derive a prediction for the time-dependent concentration of polymerized actin, which can be fitted to the fluorescence data (Fig. S5B).

We restrict ourselves to a minimal set of relevant processes: the nucleation of critical F-actin nuclei of size *γ* with a rate constant *k*_*nucl*_, and G-actin depletion due to elongation of existing F-actin filaments at an elongation rate *k*_*on*_. We reason that short filaments tend to disassemble when the free ATP-G-actin concentration *c* is below the critical concentration for elongation *C*_*c*_ = *k*_*off*_/*k*_*on*_. As a result, nucleation events in this regime are typically unproductive and need not be considered in our formalism. We thus adopt an expression for the net nucleation rate *r*_*nucl*_(*c*) that vanishes at *C* = *C*_*c*_, and satisfies the dependence at large actin concentrations consistent with F-actin nuclei of size *ϒ, r*_*nucl*_ (*c*) = *k*_*nucl*_ (*c* - *C*_*c*_)*^ϒ^*. Accordingly, we propose the following coupled time evolution of the F-actin concentration *n*(*t*) and the concentration of free G-actin in solution *c*(*t*)

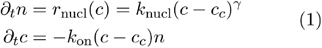

Initially, *c* is given by the total actin concentration, *c*(*t* = 0) = *C*_0_, and the F-actin concentration vanishes, *n*(*t* = 0) = 0. Due to the simplicity of this model, we can solve it analytically as detailed below.

To begin with, we nondimensionalize the coupled time evolution in Eq. (1) by introducing a typical scale of the concentration of nucleated filaments *n*_*_ and a typical time scale τ over which the monomer pool is depleted

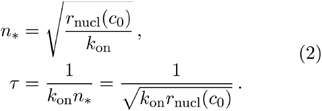

*n*_*_ is determined by the competition between nucleation *k*_*nucl*_ and elongation *k*_*on*_ for the limited monomer pool *c*_0_. The faster nucleation proceeds compared to elongation, the more actin filaments will be present in the end. Meanwhile, the time scale τ of monomer depletion is shorter the more growing filaments there are. After defining rescaled variables via 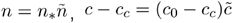 and 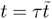, the nondimensionalized dynamics reads

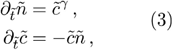

with initial conditions 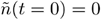 and 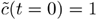. For notational convenience we drop the tildes in the following.

The time evolution for *c* can be rewritten using the natural logarithm as ∂_*t*_ ln *c* = −*n*. Solving this equation for *c*, we find

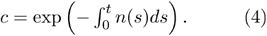

Plugging this expression into the time evolution of *n*, we obtain a differential equation in a single unknown:

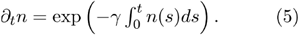

Multiplying both sides by *n* yields

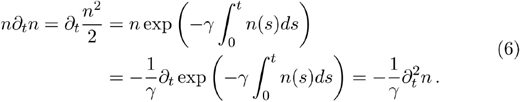

Integrating with respect to time and using Eq. (5), we arrive at a much simpler differential equation for *n*:

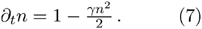

This ordinary differential equation can be solved by using separation of variables followed by partial fraction decomposition. The result reads

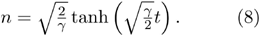

From this, we compute *c* using Eq. (4)

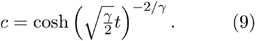

Restoring dimensions we finally obtain the result for the barbed end concentration *n*(*t*) and the G-actin concentration *c*(*t*) presented in the main text

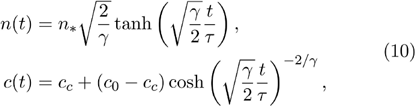

with the typical scales 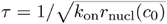 and 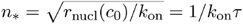. Asserting that the tetramer is the critical nucleus implies ϒ=4. As expected, *n* saturates to a value 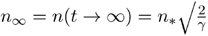, due to depletion of the monomer pool to the critical concentration, *c*_∞_ = *c*(*t* → ∞) = *c*_*c*_. From Eq. (10) we directly obtain the concentration of polymerized actin *c*_*p*_(*t*) given by

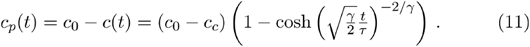

The only unknown parameter of our model is the nucleation rate constant *k*_*nucl*_, with all other parameters being fixed through independent experimental measurements. For each replicate, we determine *k*_*nucl*_ by fitting Eq. (11) to the full time course of polymerized actin concentration (Fig. S5B). Finally, this yields the total initial nucleation rate within our model

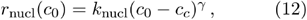

which we compare to the empirical value obtained from fitting only the early time regime in Fig. 4B.

#### Thermodynamic binding model of the actin-vimentin interaction

Here we introduce the thermodynamic model of binding for the actin-vimentin interaction, which we used to fit the saturation curves in the actin polymerization assay (Fig. 3D) and the magnetic pulldown assay (Fig. 5D). Both curves exhibit a nonlinear increase and reach saturation at concentrations much higher than that of the limiting component, namely the barbed ends (Fig. 3D) and G-actin (Fig. 5D), respectively. This indicates that both assays probe the “binding” regime (24), where the vimentin concentration needed to reach half-saturation provides an estimate for the dissociation constant. In Sec. A, we recall the binding equilibrium of ligands in a solution with fixed binding sites. We modulate this general prediction to account for the two assays used in the main text. In Sec. B we move away from the assumption of 1:1 binding stoichiometry between G-actin and vimentin monomers in the magnetic pulldown assay used in the main text. In doing so, we identify a lower bound on the number of actin binding sites per vimentin cross-section.

#### A. Thermodynamic binding model

We consider the binding thermodynamics of ligands and binding sites at 1:1 stoichiometry. The number of bound ligands is determined by the competition between the translational entropy of free ligands and the binding energy as well as the configurational entropy associated to binding. In thermodynamic equilibrium, this yields a condition relating the fraction of occupied binding sites *θ*, the total concentration of ligands *c*_0_ and of binding sites *ρ* to the dissociation constant *K*_*d*_ of a single ligand-site interaction:

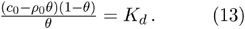

Interpreting Eq. (13) according to the experimental assay of interest allows us to probe the interactions between actin and vimentin in Figures 3D and 5D of the main text.

In the actin polymerization assay, we measured a dose-dependence of the increase in actin barbed end elongation rate on Vimv^117L^ concentration *C*_0_= [VIM^Y117L^(Fig. 3D). While we did not resolve the microscopic mechanism, we propose a putative direct interaction between actin barbed ends and VimY^117L^ ULFs as the simplest explanation. Assuming each barbed end presents a binding site to vimentin, the total concentration of binding sites *ρ*_0_ is that of the barbed ends. In this setting, the vimentin ULFs, i.e. the ligands, are available in large excess compared to the barbed ends, *c*_0_ »*ρ*_*0*_ Thus, Eq. (13) simplifies to the well-known Langmuir adsorption isotherm for the occupancy *θ* of binding sites

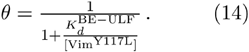

In line with our observations in Fig. 3D, this factor justifies a saturation in the actin elongation rate once the barbed end is always occupied by vimentin ULFs. Relatedly, we analyzed how the presence of vimentin in solution modulates the on- and off-rates of ATP-G-actin at the F-actin barbed end, and *K*_on_ and *k*_of_ (Fig. 1F). The dominant effect is on the off-rate, which is diminished considerably, leading to an effective increase in actin elongation rate. Notably, we measured these rates at a large vimentin concentration of 1 mg/ml, which puts us well into the plateau regime of the saturation curve in Fig. 3D. The rates measured in Fig. 1F thus probe the maximum increase in elongation rate Δ*v*_*max*_ is given by the difference in the measured elongation rate 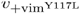 and when vimentin is present when it is absent *u*_ctrl_. We have, which

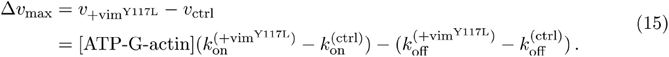

We compute this expression separately for each of the three replicates in Fig. 1F. The mean of the three resulting values and its standard error (SEM) read Δ*umax* =0.91 ±0.18s^-1^. Combining the proposed binding between vimentin ULFs and the actin barbed end, Eq. (14), with the associated stabilization of the barbed end, Eq. (15), we finally obtain the increase in actin elongation rate Δas a function of vimentin Y117L concentration

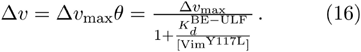

We determine the only unknown parameter, namely the apparent dissociation constant 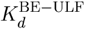 between vimentin Y117L ULFs and the F-actin barbed end, by fitting this dependence to the saturation curve in Fig. 3D of the main text.

In the magnetic pulldown assay, we measured the fraction of available G-actin *p* specifically pulled with vimentin wild-type filaments (Fig. 5D). Within our terminology, vimentin features binding sites that can be occupied by G-actin, here the ligands. In order to translate Eq. (13) to predict the fraction of bound ligands *p* instead, we use 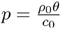 and obtain

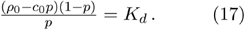

In the main text, we assumed a 1:1 stoichiometry between G-actin and vimentin monomers as a first approach to extract the apparent dissociation constant 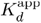. Hence, the total concentration of binding sites is given by the vimentin monomer concentration,*p0* = [Vim ^WT^]. We simplify Eq. (17) by introducing the concentration of unoccupied binding sites on vimentin 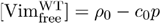. This yields the following Langmuir-like prediction for the fraction of actin *P* specifically bound to vimentin

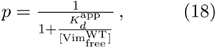

which we fit to the saturation curve in Fig. 5D to determine the only unknown parameter, the apparent dissociation constant of the interaction between ATP-G-actin and vimentin filaments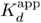.

#### B. Each vimentin cross-section has multiple binding sites for G-actin

When fitting the binding curve of the magnetic pulldown assay (Fig. 5D), we assumed a 1:1 stoichiometry between ATP-G-actin and vimentin monomers as a first simple approach, with no prior molecular information on the number of actin-binding sites per vimentin monomer, the location of these sites and how they are exposed in filamentous vimentin. The simplest situation leading to this stoichiometry would be that a vimentin monomer has one actin-binding site, which would be equally exhibited by each vimentin monomer in filamentous vimentin. However, not all binding sites might be accessible for ATP-G-actin to interact with in filamentous vimentin, especially when vimentin is attached to a magnetic bead. Here we relax the assumption of 1:1 stoichiometry between actin and vimentin monomers by introducing an additional unknown parameter, namely the number of accessible binding sites per vimentin cross-section. By studying the impact of varying this parameter on our binding model’s consistency with the experimental data, we conclude that each vimentin filament cross-section has at least 5 accessible binding sites for ATP-G-actin.

By solving Eq. (17) directly, we arrive at a more general functional form for the fraction of actin specifically bound to vimentin:

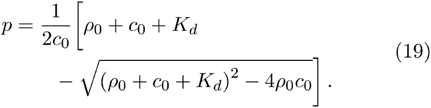

While *c*_0_ = *c*_*a*_ here simply denotes the total ATP-G-actin concentration, the distribution of binding sites on vimentin is unknown. Each vimentin cross-section is composed of *n*_*cs*_ =40 monomers (25). We denote the number of accessible binding sites per vimentin cross-section with *b*.Given the concentration of vimentin monomers *c*_*v*_, this leads to a binding site concentration of 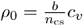. The assumption of 1:1 stoichiometry in the main text amounts to fixing *b = n_cs_* =40. In that case, Eq. (19) reduces to the simplified form in Eq. (18) such that 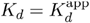. Plugging the general form *ρ*_0_ of into Eq. (19), we obtain the fraction *p* of specifically bound ATP-G-actin as a function of the total vimentin monomer concentration*c_v_*

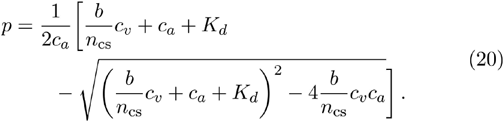

Relaxing the 1:1 stoichiometry assumption has introduced a second unknown parameter, namely the number *b* of accessible binding sites per vimentin cross-section. However, using Eq. (20) to simultaneously fit the dissociation constant *K_d_* and the number *b* of binding sites per vimentin cross-section is unfeasible: If the binding sites on vimentin are in excess,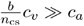, then the fraction*p* of bound ATP-G-actin is only influenced by the ratio of the two unknown parameters

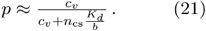

As a result, many combinations of *K*_*d*_ and *b* are consistent with the experimental data. In other terms, the fraction *p* of bound ligands has a soft mode in the parameter space. This is not surprising since many binding sites with weak interactions may bind a similar amount of G-actin as fewer binding sites with stronger interactions.

Nevertheless, we can give phenomenological bounds on both the dissociation constant *K*_*d*_ and the number of binding sites per vimentin cross-section. To this end, we proceed by elimination: We fix one of the parameters,*P*_1_, and then fit only the other one, *P*_2_, according to Eq. (20). If the resulting fit is inconsistent with our experimental data, then *P*_1_is discarded. By trying different candidates for *P*_1,_ we are able to narrow down a valid range for this parameter. The fitting results for a list of candidates for *b* and *K*_*d*_ each are shown in Fig. S6.

In Panel A, we fix *b* and fit *K*_*d*_. It illustrates that too low values for the number *b* of binding sites are inconsistent with the data. The fitted curves for *b* ϵ {1,2} are straight lines, meaning that the binding sites on vimentin are always saturated. Still, these straight lines underestimate the fraction of bound ATP-G-actin. The lowest number *b* of binding sites per vimentin cross-section consistent with the data is *b*^(min)^ ∼ 4. Relatedly, it was recently argued that the vimentin cross-section may be made up of two distinct subpopulations of five tetramers, half of which are exchangeable with the surroundings and half immobile (5). If tetramers within a subpopulation were otherwise equivalent in terms of their capability to bind ATP-G-actin, this would suggest a lower bound of *b*^(min)^ = 4, in line with our fits. Thus, our data is consistent with the hypothesis that each vimentin tetramer has at least one binding site, but leaves the question whether each tetramer within a cross-section is accessible for G-actin unanswered (5).

In Panel B we fix *K*_*d*_ and fit *b*. To bound the amount *b* of binding sites per vimentin cross-section involved in the interaction from above, we note that ATP-G-actin might only recognize vimentin oligomers instead of a motif on each individual vimentin monomer. Also, some binding sites might not be accessible due to being localized inside the filament or due to the bead. Hence, we assert that there is at most one accessible binding site per vimentin monomer, *b*^(max)^ ∼ 40. By comparing this hypothesis to the fitting results for in Panel B, we thus find an approximate upper bound for the interaction constant,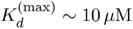, expressed in terms of vimentin monomers.

To conclude, we hypothesize a lower bound for the number of accessible binding sites per vimentin cross-section, *b*^(min)^ ∼ 4, and an upper bound for the interaction constant,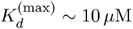, in the magnetic pulldown assay. Despite the different experimental settings in the actin polymerization assay and the magnetic pulldown assay, these bounds show that both assays may probe the same molecular interaction between actin and vimentin. Nevertheless, the exact number of accessible binding sites per vimentin cross-section remains unknown, which makes it unfeasible to compare the two assays in the main text quantitatively.

